# Comparative ACE2 variation and primate COVID-19 risk

**DOI:** 10.1101/2020.04.09.034967

**Authors:** Amanda D. Melin, Mareike C. Janiak, Frank Marrone, Paramjit S. Arora, James P. Higham

**Author notes:** These authors contributed equally to this work (Amanda D. Melin, Mareike C. Janiak). Authors for correspondence: Dr Amanda Melin, Department of Anthropology and Archaeology, University of Calgary, 2500 University Drive, Calgary, Alberta T2N 1N4, CA, Dr James Higham, Department of Anthropology, New York University, 25 Waverly Place, New York, NY 10003, US.

## Abstract

The emergence of the novel coronavirus SARS-CoV-2, which in humans is highly infectious and leads to the potentially fatal disease COVID-19, has caused hundreds of thousands of deaths and huge global disruption. The viral infection may also represent an existential threat to our closest living relatives, the nonhuman primates, many of which are endangered and often reduced to small populations. The virus engages the host cell receptor, angiotensin-converting enzyme-2 (ACE2), through the receptor binding domain (RBD) on the spike protein. The contact surface of ACE2 displays amino acid residues that are critical for virus recognition, and variations at these critical residues are likely to modulate infection susceptibility across species. While infection studies are emerging and have shown that some primates, such as rhesus macaques and vervet monkeys, develop COVID-19-like symptoms when exposed to the virus, the susceptibility of many other nonhuman primates is unknown. Here, we show that all apes, including chimpanzees, bonobos, gorillas, and orangutans, and all African and Asian monkeys (catarrhines), exhibit the same set of twelve key amino acid residues as human ACE2. Monkeys in the Americas, and some tarsiers, lemurs and lorisoids, differ at significant contact residues, and protein modeling predicts that these differences should greatly reduce the binding affinity of the ACE2 for the virus, hence moderating their susceptibility for infection. Other lemurs are predicted to be closer to catarrhines in their susceptibility. Our study suggests that apes and African and Asian monkeys, as well as some lemurs are all likely to be highly susceptible to SARS-CoV-2, representing a critical threat to their survival. Urgent actions have been undertaken to limit the exposure of Great Apes to humans, and similar efforts may be necessary for many other primate species.

## Introduction

In late 2019 a novel coronavirus SARS-CoV-2 emerged in China. In humans, this virus can lead to the respiratory disease COVID-19, which can be fatal^1,2^. Since then, SARS-CoV-2 has spread around the world, causing widespread mortality, and with major impacts on societies and economies. While the virus and its resulting disease represent a major humanitarian disaster, they also represent a potential existential risk to our closest living relatives, the nonhuman primates. Transmission incidences of bacteria and viruses - including another coronavirus (H-CoV-OC43) - from humans to wild populations of nonhuman primates have previously caused outbreaks of Ebola, yellow fever, and fatal respiratory diseases, leading in some cases to mass mortality^3–9^. Such past events raise considerable concerns among the global conservation community with respect to the impact of the current pandemic^10^.

Infection studies of rhesus monkeys, longtailed macaques, and vervets as biomedical models have made it clear that at least some nonhuman primate species are permissive to SARS-CoV-2 infection and develop symptoms in response to infection that resemble those of humans following the development of COVID-19, including similar age-related effects^11–16^. Recognizing the potential danger of COVID-19 to nonhuman primates, the International Union for the Conservation of Nature (IUCN), together with the Great Apes section of the Primate Specialist Group, released a joint statement on precautions that should be taken for researchers and caretakers when interacting with great apes^17^. However, the risk for many primate taxa remains unknown. Here we begin to assess the potential likelihood that our closest living relatives are susceptible to SARS-CoV-2 infection.

While the biology underlying susceptibility to SARS-CoV-2 infection remains to be fully elucidated, the viral target is well established. The SARS-CoV-2 virus binds to the cellulSar receptor protein angiotensin-converting enzyme-2 (ACE2), which is expressed on the extracellular surface of endothelial cells of diverse bodily tissues, including the lungs, kidneys, small intestine and renal tubes^18^. ACE2 is a carboxypeptidase whose activities include regulation of blood pressure and inflammatory response through its role in cleaving the vasoconstrictor angiotensin II to produce angiotensin 1-7 and triggering varied downstream responses^19–22^. ACE2 is made up of a signal sequence at the N-terminus (residues 1-17), a transmembrane sequence at the C-terminus (residues 741-762), and an extracellular region, which contains a zinc metallopeptidase domain (residues 19–611) and a collectrin homolog (residues 612-740)^23,24^.

Characterizations of the infection dynamics of SARS-CoV-2 have demonstrated that the binding affinity for the human ACE2 receptor is high, which is a key factor in determining the susceptibility and transmission dynamics. When compared to SARS-CoV, which caused a serious global outbreak of disease in 2002-2003^25,26^, the binding affinity between SARS-CoV2 and ACE2 is estimated to be between 4-fold^27–30^ and 10- to 20-fold greater^31^. Recent reports describing structural characterization of ACE2 in complex with the SARS-CoV2 spike protein receptor binding domain (RBD)^27–30^ allow identification of the key binding residues that enable the host-pathogen protein-protein recognition. Following initial binding of the virus to the ACE2 receptor, humans experience a great deal of variation in response to infection, with some individuals experiencing relatively mild symptoms, while others experience major breathing problems and organ failures, which can lead to death. Some of this response is known to be linked to variation in how the immune system responds to infection, with some individuals experiencing a hyperinflammatory ‘cytokine storm’, which in turn aggravates respiratory failures and increases mortality risk^32,33^. There may also be some variation among humans in initial susceptibility to infection, such that approaches examining variation in ACE2 tissue expression and gene sequences can offer insight into variation in human susceptibility to COVID-19^34–37^. Similarly, we can use such an approach to compare sequence variation across species, and hence try to predict the likely interspecific variation in susceptibility to initial infection. Previous analysis of comparative variation at these sites enabled estimates of the affinity of the ACE2 receptor for SARS-CoV in nonhuman species (bats)^38^.

Here, we undertake such an analysis for SARS-CoV-2 across the primate radiation. Our aim is to investigate the likelihood of initial susceptibility to infection for different major radiations and species, while recognizing that down-stream processes such as immune responses are likely to determine the extent to which species and individuals develop symptoms and pathologies in response to infection. We compiled *ACE2* gene sequence data from 29 primate species for which genomes are publicly available, covering primate taxonomic breadth. For comparison, we assessed 4 species of other mammals that have been tested directly for SARS-CoV2 susceptibility in laboratory infection studies^39^. We also included in our analysis the amino acid sequence variation at these sites for horseshoe bats, thought to be the original vector of the virus, and pangolins, a potential intermediate host, where viral recombination may have led to the novel viral form SARS-CoV-2^40^. We assessed the variation at amino acid residues identified as critical for ACE2 recognition by CoV RBD, and undertook analysis of positive selection and protein modeling to gauge the potential for adaptive differences and the likely effects of protein variation. Our aim was to develop predictions about the susceptibility of our closest living relatives to SARS-CoV-2 as a resource for stakeholders, including researchers, caretakers, practitioners, conservationists, and governmental and nongovernmental agencies.

## Methods

### Variation in ACE2 sequences

We compiled *ACE2* gene sequences for 16 catarrhine primates: 4 species from all 3 genera of great ape (*Gorilla*, *Pan*, *Pongo*), 2 genera of gibbons (*Hylobates*, *Nomascus*), and 10 species of African and Asian monkeys in 7 genera (*Cercocebus, Chlorocebus, Macaca, Mandrillus, Papio, Rhinopithecus, Piliocolobus, Theropithecus*); 6 genera of platyrrhines (monkeys from the Americas: *Alouatta, Aotus, Callithrix, Cebus, Saimiri, Sapajus*); 1 species of tarsier (*Carlito syrichta)*; and 5 genera of strepsirrhines (lemurs and lorisoids: *Eulemur*, *Daubentonia*, *Microcebus*, *Propithecus*, *Otolemur*) (Suppl. Table S1). We also included 4 species of mammals that have been tested clinically for susceptibility to SARS-CoV-2 infection^39^, including the domestic cat (*Felis catus*), dog (*Canis lupus familiaris*), pig (*Sus scrofa*), and ferret (*Mustela putorius furo*). Finally, we included the pangolin (*Manis javanica*) and several bat species, including horseshoe bats (*Rhinolophus* spp., *Hipposideros pratti*, *Myotis daubentonii*). Sequences were retrieved from NCBI, either from annotations of published genomes or from GenBank entries^38^. We manually checked annotations by performing tblastn searches of the human ACE2 protein sequence against each genome. We identified one misannotation for exon 15 in *Microcebus murinus*, which we manually corrected. The *ACE2* nucleotide sequence for *Alouatta palliata* was obtained from an unpublished draft genome, via tblastn searches using the *Cebus capucinus* ACE2 protein sequence as a query and default search settings. Accession numbers for sequences retrieved from NCBI and GenBank are provided in Supplemental Table S1 and the *Alouatta palliata* sequence is available in the supplemental materials.

Coding sequences were translated using Geneious Version 9.1.8 and we aligned both nucleotide and amino acid sequences with MAFFT^41^. Amino acids were aligned with the BLOSUM62 scoring matrix, while the 200 PAM scoring matrix was used for nucleotides. A 1.53 gap open penalty and an offset value of 0.123 were used for both. We manually inspected and corrected any misalignments, and verified the absence of indels and premature stop codons.

To visualize patterns of gene conservation across taxa and identify the congruence of the *ACE2* gene tree with currently accepted phylogenetic relationships among species, we reconstructed trees using both Bayesian (MrBayes 3.2.6^42^) and Maximum Likelihood (RAxML 8.2.11^43^) methods with 200,000 MCMC cycles and 1,000 bootstrap replicates, respectively (code available on GitHub^44^). Gene trees were compared to a current species phylogeny assembled using TimeTree^45^, which is also used to illustrate the evolutionary relationships between study species in Figure 1. Parsimony-informative sites along the *ACE2* sequence were identified with the pis function in the R package ips v. 0.0.11^46,47^.

**Figure 1.**
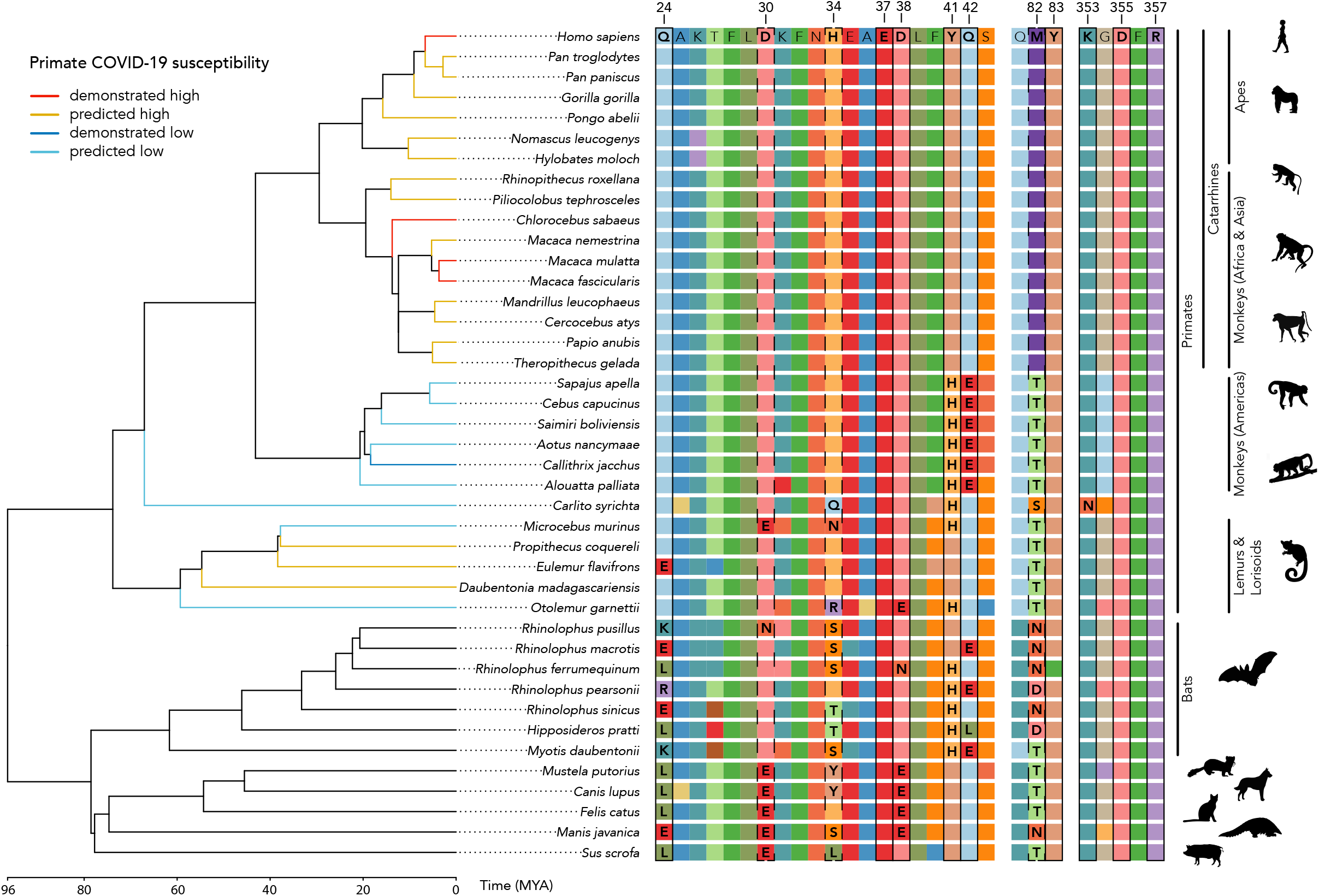
ACE2 protein sequence alignment and evolutionary relationships of study species. Branch lengths represent evolutionary distance (time, in millions of years) estimated from TimeTree^45^. We outline amino acid residues at critical binding sites for the SARS-CoV-2 spike receptor binding domain. Solid outlines highlight sites predicted to have the most substantial impact on viral binding affinity. Notably, protein sequences of catarrhine primates are highly conserved, including uniformity among amino acids at all binding sites. Primate species that are able to be successfully infected with COVID-19 are indicated in red. Predicted susceptibility to COVID-19 for other primates is additionally coded by terminal branch colors.

### Identification of critical binding residues and species-specific ACE2–RBD interactions

Critical ACE2 protein contact sites for the viral spike protein receptor binding domain (RBD) have been identified using cryo EM and X-ray crystallography structural analysis methods^27–30^. The ACE2-RBD complex is characteristic of protein-protein interactions (PPIs) that feature extended interfaces spanning a multitude of binding residues. Experimental and computational analyses of PPIs have shown that a handful of contact residues can dominate the binding energy landscape^48^. Alanine scanning mutagenesis provides an assessment of the contribution of each residue to complex formation^49–51^. Critical binding residues can be computationally identified by assessing the change in binding free energy of complex formation upon mutation of the particular residue to alanine, which is the smallest residue that may be incorporated without significantly impacting the protein backbone conformation^52^. Our computational modeling utilizes the human SARS RBD/ACE2 high resolution structures, and we make the implicit assumption that the overall conformation of ACE2 is conserved among different species. This assumption, which is rooted in the high sequence similarity between ACE2 sequences, allows us to use the structure of the complex to predict the impact of mutations at the protein-protein interface.

We defined critical residues as those that upon mutation to alanine decrease the binding energy by a threshold value ΔΔG_bind_ ≥1.0 kcal/mol. Nine of the 21 residues identified by alanine scanning as involved in the ACE2-RBD complex met this criterion (Suppl. Table S2). There was large congruence in the sites identified with those highlighted by other methods. Each of the eight sites implicated by cryo EM^27^, were also detected by alanine modelling; five residues were ≥1.0 kcal/mol threshold and 3 were below this threshold. To be cautious, in addition to the the 9 critical ACE2 sites we identified through alanine scanning, we also examined residue variation at the 3 sites that fell below the ≥1.0 kcal/mol threshold but that were identified as important by structural analyses^27–30^ for a total of 12 critical sites. All computational alanine scanning mutagenesis analyses were performed using Rosetta software^52^. The alanine mutagenesis approach has been extensively evaluated and used to analyze PPIs and design their inhibitors, including by members of the present authorship^53,54^.

We utilized the SSIPe program^55^ to predict how ACE2 amino acid differences in each species would affect the relative binding energy of the ACE2/SARS-Cov-2 interaction. Using human ACE2 bound to the SARS-Cov-2 RBD as a benchmark (PDB 6M0J), the program mutates selected residues and compares the binding energy to that of the original. Using this algorithm, we studied interactions of all primates across the full suite of amino acid changes occurring at critical binding sites for each species. To more thoroughly assess the impact of each amino acid substitution, we also examined the predicted effect of individual amino acid changes (in isolation) on protein-binding affinity.

### Adaptive evolution of ACE2 sequences

We further investigated *ACE2* and how selective pressures in different clades might be shaping variation at the binding sites, using codeml clade C and branch-site models in PAML^56^. We first tested if selection acting on *ACE2* is divergent between the major clades in our sample (platyrrhine, catarrhine, and strepsirrhine primates, non-primate mammals) with the codeml clade model C, which was compared to the null model (M2a_rel) with a likelihood ratio test^57^. This test shows whether there is divergent selection (dN/dS ratio = *ω*) across all clades, but not which clades are experiencing positive selection. We, therefore, followed the clade model with a series of branch-site models, which allow one clade at a time to be designated as a set of “foreground” branches and test whether this clade has experienced episodes of positive selection compared to the remaining sets of “background” branches (*ω*_foreground_ > *ω*_background_). Branch-site models are compared to a null model that fixes *ω* at 1 with a likelihood ratio test. In the case of the alternative model having a significantly better fit than the null model, indicating positive selection, potential sites under positive selection are identified with a Bayes Empirical Bayes (BEB) approach^58^. We completed branch-site models for each primate clade (platyrrhine, strepsirrhine, and catarrhine), as well as bats, because previous research has identified *ACE2* to be under positive selection in this clade, potentially in response to coronaviruses^59^. We had to exclude *Hipposideros pratti* and *Myotis daubentonii* from PAML analyses, because only a partial *ACE2* sequence was available for these two species. Input files and control files for PAML codeml analyses are available in the GitHub repository^44^.

## Results

### Variation in ACE2 sequences

The *ACE2* gene (2418 bp) and translated protein (805 amino acids) sequences are strongly conserved across primates. The average pairwise identity across 29 primate species is 93.6% for the *ACE2* nucleotide sequence and 90.8% for the protein sequence, with a pairwise similarity (BLOSUM62 ≥ 1) of 95.3% (Suppl. Tables S3-5). Out of 2418 bp, 1631 bp (67.5%) are identical, while 401 bp (16.58%) are phylogenetically informative sites for primates, and gene trees we generated (Suppl. Fig. S1a,b) closely recapitulate the currently accepted phylogeny of primates (Figure 1). In particular, the twelve sites in the ACE2 protein that are critical for binding of the SARS-CoV-2 virus are invariant across the Catarrhini, which includes great apes, gibbons, and monkeys of Africa and Asia (Figure 1). Furthermore, catarrhines do not vary at any of the 21 sites identified by alanine scanning (Suppl. Table S2, Suppl. Fig. S2). The other major radiation of monkeys, those found in the Americas (Platyrrhini), have ACE2 sequences that are less similar to humans across the length of the protein (91.68-92.55% identical to *H. sapiens*, Suppl. Table S4) but conserved within their clade (average pairwise identity 97.2%, Suppl. Table S4). They share nine of twelve critical amino acid residues with catarrhine primates; the three sites that vary from catarrhines, H41, E42 and T82, are conserved within the platyrrhines. Strepsirrhine primates and tarsiers, were more variable in the binding sites and less similar to the human protein across the length of the sequence (81.86-86.93% pairwise identity, Suppl. Table S4). Like platyrrhines, the tarsier (*Carlito syrichta*), mouse lemur *(Microcebus murinus)*, and galago (*Otolemur garnettii*) have an H41 residue, while the sifaka (*Propithecus coquereli*), aye-aye (*Daubentonia madagascariensis*), and the blue-eyed black lemur (*Eulemur flavifrons*) have the same allele as humans and other catarrhines, Y41.

In non-primate mammals, a higher number of amino acid substitutions are evident (77.37-85.22% pairwise identity to *H. sapiens*, Suppl. Table S4), including at critical binding sites. All species possess a different residue to primates at site 24. Bats are exceptionally variable within the binding sites, with the genus *Rhinolophus* alone encompassing all of the variation seen in the rest of the non-primate mammals. Where primates have glutamine (Q24), bats have glutamate (E24), lysine (K24), leucine (L24), or arginine (R24) (Figure 1). All fasta alignments of ACE2 gene and protein sequences are available in the supplemental materials, a full-length protein alignment is shown in Suppl. Figure S2, and distance matrices are provided in Suppl. Table S3-5.

### Analysis of species-specific residues on ACE2–RBD interactions

The ACE2 receptors of all catarrhines have identical residues to humans at the RBD/ACE2 binding interface across all 12 critical sites and are predicted to have similar binding affinity for SARS-CoV-2. Platyrrhines diverge from catarrhines at three of the twelve critical amino acid residues. Compared to catarrhine ACE2, the platyrrhines’ ACE2 is predicted to bind SARS-CoV2 RBD with roughly 400-fold reduced affinity (ΔΔG_bind_=3.5 kcal/mol) (Table 1a). In particular, the change at site 41 from Y to H found in monkeys in the Americas has the largest impact of any residue change examined (Table 1b), which alone is predicted to lead to a 25-fold decrease in the binding affinity to SARS-CoV-2 (Figure 2). This single mutation combined with additional substitutions, especially Q42E, found in platyrrhines is predicted to significantly reduce the likelihood of successful viral binding (Table 1b). Of the other primates modeled, two of the three strepsirhines, and tarsiers, also have the H41 residue and furthermore have additional protein sequence differences leading to further decreases in predicted binding affinity. The predicted binding affinity of tarsier ACE2 is the most dissimilar to humans and this primate might be the least susceptible of the species we examine. In contrast, Coquerel’s sifaka (*Propithecus coquereli*), the aye-aye (*Daubentonia madagascariensis*), and blue-eyed black lemur (*Eulemur flavifrons*) share the same residue as humans and other catarrhines at site 41 and have projected affinities that are near to humans (Table 1b). Other mammals included in our study - ferrets, cats, dogs, pigs, pangolin and two of the seven bat species (*R. pusillus* and *R. macrotis)* - show the same residue as humans (Y) at site 41, with accompanying strong affinities for SARS-CoV-2. The remaining five sister species of bats possess H41 and lower binding affinities (Table 1b).

**Table 1.** Results of computational protein-protein interaction experiments predicting impact of amino acid changes, relative to human ACE2 residues, at critical binding sites with SARS-CoV-2 receptor binding domain. Impacts of changes across the full complement of critical binding sites are presented in (A), single residue replacements are presented in (B).

| Species | Mutations | $\Delta\Delta G$ (kcal/mol)<sup>a</sup> |
| --- | --- | --- |
| *Carlito syrichta* | H34Q, Y41H, M82S, K353N | 5.506 |
| *Microcebus murinus* | D30E, H34N, Y41H, M82T | 4.001 |
| *Propithecus coquereli* | M82T | 0.938 |
| *Otolemur garnettii* | H34R, D38E, Y41H, M82T | 3.815 |
| Monkeys (Americas) | Y41H, Q42E, M42T | 3.506 |

**B)**
| Mutation | $\Delta\Delta G$ (kcal/mol)<sup>a</sup> |
| --- | --- |
| Y41H | 1.929 |
| Q42E | 0.954 |
| M82T | 0.938 |
| D38E | 0.651 |
| Q24L | -0.753 |
| H34L | -0.566 |
| H34Y | -0.139 |
| D30E | 0.692 |

<sup>a</sup>Mutations were analyzed with SSIPe server (https://zhanglab.ccmb.med.umich.edu/SSIPe/) and PDB file 6M0J.

**Figure 2.**
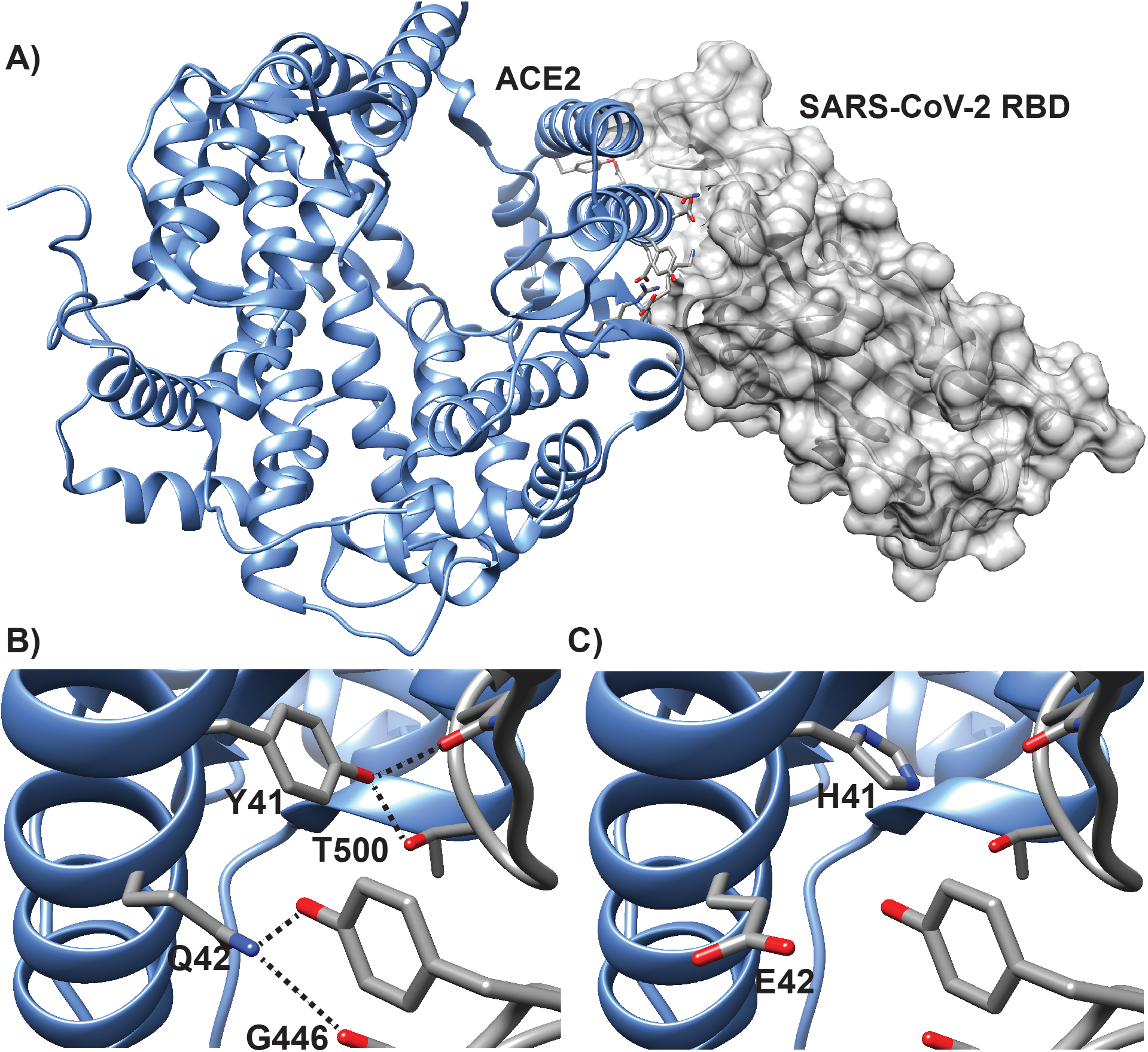
Model of human ACE2 in complex with SARS-CoV-2 RBD. Key ACE2 interfacial residues are highlighted. (A). Interactions at critical binding sites 41 and 42 are shown for the residues found in all catarrhines (apes and monkeys in Africa and Asia); (B), and for the residues found in all platyrrhines (monkeys in the Americas) (C). The dashed lines indicate predicted hydrogen bonding interactions. Y41 participates in extensive van der Waals and hydrogen bonding interactions with RBD; these interactions are abrogated with histidine. Q42 side chain amide serves as a hydrogen acceptor and donor to contact RBD; change to glutamic acid diminishes the hydrogen bonding interactions.

### Adaptive evolution of ACE2 sequences

We find evidence that the selective pressures acting on *ACE2* are not equivalent across the major clades in our analysis. The codeml clade model C provided a better fit than the null model (LRT = 26.726, p < 0.001; Table 2, Suppl. Table S6). Branch-site models indicate that the catarrhine primate clade (LRT = 14.546, p < 0.001) and bat clade (LRT = 42.649, p < 0.001) are both under positive selection, while platyrrhines (LRT = 0.633, p = 0.427) and strepsirrhines (LRT = 0.833, p = 0.361) are not. The six positively selected sites in the bat clade include the binding site 24 and two others adjacent to known binding sites (Table 2). In catarrhines, the three positively selected sites identified by BEB calculations are not near the binding sites for SARS-Cov-2 (residues 249, 653, and 658; Table 2).

**Table 2.** Results of codeml analyses of adaptive evolution across ACE2 gene sequences.

| Model | Foreground Branch | $\omega$ | proportion of sites | LRT | $p$ | positively selected sites<sup>a,b</sup> |
| --- | --- | --- | --- | --- | --- | --- |
| clade C | n/a | $\omega_0 = 0.059$,<br>$\omega_1 = 1.000$,<br>$\omega_2 = 0.081$,<br>$\omega_3 = 1.123$,<br>$\omega_4 = 0.236$,<br>$\omega_5 = 1.346$ | $p_0 = 0.581$,<br>$p_1 = 0.331$,<br>$p_{2-5} = 0.089$ | 26.726 | <0.001 | n/a |
| branch-site | platyrrhines | background:<br>$\omega_0 = 0.076$,<br>$\omega_1 = 1.000$,<br>$\omega_{2a} = 0.076$,<br>$\omega_{2b} = 1.000$;<br>foreground:<br>$\omega_0 = 0.076$,<br>$\omega_1 = 1.000$,<br>$\omega_{2a} = 6.218$,<br>$\omega_{2b} = 6.218$ | $p_0 = 0.638$,<br>$p_1 = 0.359$,<br>$p_{2a} = 0.002$,<br>$p_{2b} = 0.001$ | 0.633 | 0.427 | none |
| | catarrhines | background:<br>$\omega_0 = 0.075$,<br>$\omega_1 = 1.000$,<br>$\omega_{2a} = 0.075$,<br>$\omega_{2b} = 1.000$;<br>foreground:<br>$\omega_0 = 0.075$,<br>$\omega_1 = 1.000$,<br>$\omega_{2a} = 8.988$,<br>$\omega_{2b} = 8.988$ | $p_0 = 0.631$,<br>$p_1 = 0.356$,<br>$p_{2a} = 0.009$,<br>$p_{2b} = 0.005$ | 14.546 | 0.0001 | 249M (0.962*), 653A (0.958*), 658V (0.957*) |
| | strepsirrhines | background:<br>$\omega_0 = 0.072$,<br>$\omega_1 = 1.000$,<br>$\omega_{2a} = 0.072$,<br>$\omega_{2b} = 1.000$;<br>foreground:<br>$\omega_0 = 0.072$,<br>$\omega_1 = 1.000$,<br>$\omega_{2a} = 1.384$,<br>$\omega_{2b} = 1.384$ | $p_0 = 0.607$,<br>$p_1 = 0.316$,<br>$p_{2a} = 0.051$,<br>$p_{2b} = 0.027$ | 0.833 | 0.361 | none |
| | bats | background:<br>$\omega_0 = 0.075$,<br>$\omega_1 = 1.000$,<br>$\omega_{2a} = 0.075$,<br>$\omega_{2b} = 1.000$;<br>foreground:<br>$\omega_0 = 0.075$,<br>$\omega_1 = 1.000$,<br>$\omega_{2a} = 10.535$,<br>$\omega_{2b} = 10.535$ | $p_0 = 0.626$,<br>$p_1 = 0.338$,<br>$p_{2a} = 0.024$,<br>$p_{2b} = 0.013$ | 42.649 | <0.001 | 24Q (0.998**), 31E (0.959*), 35E (0.974*), 298V (0.959*), 568L (0.998**), 575G (0.965*) |

<sup>a</sup>Sites with posterior probability >0.95 are shown. <sup>b</sup>The amino acid shown reflects the residue present at the site in the first sequence of the alignment (*Alouatta palliata*).

## Discussion

Our results strongly suggest that catarrhines - all apes, and all monkeys of Africa and Asia, are likely to be susceptible to infection by SARS-CoV-2. There is high conservancy in the protein sequence of the target receptor, ACE2, including uniformity at all identified and tested major binding sites. Indeed, even among the 21 residues identified in our full list of potential binding points, catarrhines are invariant (Suppl. Table 2, Suppl. Fig. S2). Consistent with our results, infection studies show that rhesus monkeys (*Macaca mulatta*), longtailed macaques (*M. fascicularis*) and vervets (*Chlorocebus sabaeus*) are permissive to infection by SARS-CoV-2, and go on to develop COVID-19 like symptoms^11–14,16^. Our results based on protein modeling offer potentially better news for monkeys in the Americas (platyrrhines). There are three differences in amino acid residues between platyrrhines and catarrhines, and two of these, H41Y and E42Q show strong evidence of being impactful changes. These amino acid changes are modeled to reduce the binding affinity between SARS-CoV-2 and ACE2 by ca. 400-fold. Recent clinical analysis of viral shedding, viremia, and histopathology in catarrhine (macaque) versus platyrrhine (marmoset, *Callithrix jacchus*) responses to inoculation with SARS-CoV-2, show much more severe presentation of disease symptoms in the former, strongly supporting our results^13^. Similar reduced susceptibility is predicted for tarsiers, and two of the five lemurs and lorisoids (strepsirrhines). What is concerning is three of the analyzed lemurs spanning divergent lineages - the Coquerel’s sifaka, the aye-aye, and the blue-eyed black lemur - are more similar to catarrhines at important binding sites, including possessing the high risk residue variant at site 41, and as such are also predicted to be susceptible. Nonetheless, these are only predicted results based on amino acid residues, and protein-protein interaction models. We urge extreme caution in using our analyses as the basis for relaxing policies regarding the protection of platyrrhines, tarsiers or any strepsirrhines. Experimental assessment of synthetic protein interactions can now occur in the laboratory e.g.^60^, and confirmation of our model predictions should be sought before any firm conclusions are reached.

Emerging evidence in experimental mammalian models appears to support our results; dogs, ferrets, pigs, and cats have all shown some susceptibility to SARS-CoV-2 but have demonstrated variation in disease severity and presentation, including across studies^39,61^. Substitutions at binding sites might be at least partially protective against COVID-19 in these mammals. For example, the limited experimental evidence to date suggests that while cats - which have the same residue as humans at site 34 - are not strongly symptomatic, they present lung lesions, while dogs - which have a substitution at this site - do not^39^. The amino acid residue at site 24 differs from primates in all other mammalian species examined. However, our models suggest that the variant residues may confer relatively minor reductions in binding affinity. Other sources of variation may affect ACE2 protein stability^35^. Our results are also consistent with previous reports that ACE2 genetic diversity is greater among bats than that observed among mammals susceptible to SARS-CoV-type viruses. This variation has been suggested to indicate that bat species may act as a reservoir of SARS-CoV viruses or their progenitors^38^. Intriguingly, all but 2 bat species we examined have the putatively protective variant, H41. Additionally, results of our codeml branch-site analysis support previous findings of *ACE2* in bats being under positive selection, including sites within the binding domain of SARS-CoV and SARS-CoV-2^59^, which may be evidence of host-virus coevolution. Sites showing evidence of positive selection within catarrhine *ACE2* sequences were not in or near known CoV binding sites (Table 2, Figure 1). Two (residues 653, 658) fall within the cleavage site (residues 652-659) utilized by the sheddase ADAM17, known to interact with ACE2^62^. However, neither of the residues under selection are the amino acids targeted by ADAM17^63^ leaving the functional significance of evolution at these sites uncertain. Further clinical and laboratory study is needed to fully understand infection dynamics.

There are a number of important caveats to our study. Firstly, all of our predictions are based on interpretations of gene and resultant amino acid sequences, rather than based on direct assessment of individual responses to induced infection. Nonetheless, the overall pattern of our results is being borne out by infection studies on a few species that are used as biomedical models. So far, all catarrhine species tested by infection studies, including rhesus macaques, longtailed macaques, and vervet monkeys^13,14,64^ have exhibited COVID-19-like symptoms in response to infection, including large lung and other organ lesions^13^ and cytokine storms^14^. In contrast, marmosets did not exhibit major symptoms in response to infection^13^. While these results support and validate our findings based on ACE2 sequence interpretation, the number of primate species that can and will be tested directly by infection studies will be restricted to just a handful. Our study enhances this picture, by allowing inferences to be made across the primate radiation, backed up by the published infection studies on a few target model species.

Some of our results, such as the uniform conservation of *ACE2* among catarrhines, backed up by the demonstrated high susceptibility of humans and other catarrhines to SARS-CoV-2, should give a good degree of confidence of high levels of risk.

Given the identical residues of humans to other apes and monkeys in Asia and Africa at the target site, it seems unlikely that the ACE2 receptor and the SARS-CoV-2 proteins would not readily bind. Our results for other taxa are dependent on modeling, hence should be treated more cautiously. This includes all interpretations of the susceptibility of platyrrhines and strepsirrhines, where the effects of residue differences on binding affinities have been estimated based on protein-protein interaction modeling. Another caveat is that we have modeled only interactions at binding sites, and not predictions based on full residue sequence variation. Residues that are not in direct contact may still affect binding allosterically. Other factors, including proteases necessary for viral entry, and other viral targets, may also impact disease susceptibility and responses^35^. More generally, if adhering to the precautionary principle, then our results highlighting higher risks to some species should be taken with greater gravity than our results that predict potential lower risks to others. Another limitation of our study is that we have looked at only 29 primate species, albeit with broad taxonomic scope. Analysis of additional species is important, especially among strepsirrhine species, where our coverage is relatively scant. In particular, the residue overlap at important binding sites in the sequences of Coquerel’s sifaka, the aye-aye, and blue-eyed black lemur with those of catarrhines suggests many lemurs may be highly vulnerable and we underscore the need to assess a wider diversity of lemur species. Furthermore, we examine only one individual per species, and intraspecific variation across populations should be considered; however, studies on intraspecific ACE2 variation with humans and vervet monkeys suggest ACE2 variants are low in frequency^65–67^. Finally, it is also important to remember that our study assesses only the potential for initial binding of the virus to the target site. Downstream consequences of infection may differ drastically based on species-specific proteases, genomic variants, metabolism, and immune system responses^68,69^. In humans, the development of COVID-19 can lead to a pro-inflammatory cytokine storm of hyperinflammation, which may lead to some of the more severe impacts of infection^32,70^. Nonetheless, it is evidence from the hundreds of thousands of deaths and global lockdown that humans are highly susceptible to SARS-CoV-2 infection, and our results suggest that all apes and monkeys in Africa and Asia are similarly susceptible.

Many endangered primate species are now only found in very small population sizes^71^. For example, there are believed to be only around 1000 mountain gorillas left in their entire range^72^. With such small populations, the introduction of a new highly infectious disease is a potential extinction-level event. Re-opening access to habituated great ape groups for tourism purposes, which may be critical to local economies^73^, may be fraught with issues. IUCN best practices recommend that tourists stay at least 7 metres away from great apes^74^, but in practice, almost all tourists get far closer than this - for example, the average distance that tourists get from mountain gorillas at the Bwindi Impenetrable National Park in Uganda is just 2.76 metres^75^. Concerted effort may be required by all stakeholders to try to avoid the introduction of SARS-CoV-2 into wild primate populations^10^. Recent measures suggested by the IUCN for researchers and caretakers of great ape populations include: ensuring that all individuals wear clean clothing and disinfected footwear; providing hand-washing facilities; requiring that a surgical face mask be worn by anyone coming within 10 meters of great apes; ensuring that individuals needing to cough or sneeze ideally leave the area, or at least cough/sneeze into the crux of their elbows; imposing a 14-day quarantine for all people arriving into great ape areas who will come into frequent close proximity with them^17^. The IUCN’s ‘Best Practice Guidelines for Health Monitoring and Disease Control in Great Ape Populations’ should also be followed^76^.

Our results suggest that dozens of nonhuman primate species, including all of our closest relatives, are likely to be highly susceptible to SARS-CoV-2 infection, and vulnerable to its effects. Major actions may be needed to limit the exposure of many wild primate populations to humans. This is likely to require coordinated input from all stakeholders, including local communities, international and national governmental agencies, nongovernmental conservation and development organizations, and academics and researchers. While the focus of many at this time is rightly on mitigating the humanitarian devastation of COVID-19, we also have a duty to ensure that our closest living relatives do not suffer extinctions, or massive population declines, in response to yet another human-induced catastrophe.

## Supporting information

nucleotide sequence alignment

nucleotide sequences

protein sequence alignment

protein sequences

## Data Availability Statement

Nucleotide and protein sequences used in this study are available from NCBI and are also available as fasta files and alignments in the supplemental material and on Github (https://github.com/MareikeJaniak/ACE2). All code used in this project is available in the same repository.

## Acknowledgements

MCJ was funded by a National Sciences and Engineering Council of Canada Discovery Accelerator Supplement to ADM and by a postdoctoral fellowship from the Alberta Children’s Hospital Research Institute. PSA thanks the National Institutes of Health (R35GM130333) for financial support. We thank four reviewers for constructive comments, which improved the manuscript considerably.

## Author Contributions

ADM, JPH, and MCJ designed the study. ADM and JPH wrote the paper with input and edits from MCJ and PSA. MCJ conducted genetic analyses with input from ADM. FM ran the protein substitution models with input from PSA. All authors have approved the final submission for publication.

