## Supplemental Tables and Figures for "Comparative ACE2 variation and primate COVID-19 risk"

**Supplemental Table S1.** Gene IDs and NCBI accession numbers of *ACE2* gene sequences included in the study.

| Species | Common Name | Gene ID | NCBI Accession Number | Notes |
|---|---|---|---|---|
| *Homo sapiens* | human | 59272 | NM_001371415.1 | |
| *Pan troglodytes* | chimpanzee | 465511 | XM_016942979.1 | |
| *Pan paniscus* | bonobo | 100970340 | XM_008974180.1 | |
| *Gorilla gorilla* | gorilla | 101142534 | XM_019019204.1 | |
| *Pongo abelii* | Sumatran orangutan | 100171441 | NM_001131132.2 | |
| *Nomascus leucogenys* | northern white-cheeked gibbon | 100602708 | XM_003261084.3 | |
| *Hylobates moloch* | silvery gibbon | 116811532 | XM_032756617.1 | |
| *Rhinopithecus roxellana* | golden snub-nosed monkey | 104664530 | XM_010366065.2 | |
| *Piliocolobus tephrosceles* | Ugandan red colobus | 111531712 | XM_023199053.2 | |
| *Macaca mulatta* | rhesus macaque | 712790 | NM_001135696.1 | |
| *Macaca nemestrina* | pigtail macaque | 105478157 | XM_011735203.2 | |
| *Macaca fascicularis* | long-tailed macaque | 102130864 | XM_005593037.2 | |
| *Cercocebus atys* | sooty mangabey | 105574684 | XM_012035809.1 | |
| *Mandrillus leucophaeus* | drill | 105550583 | XM_011995533.1 | |
| *Papio anubis* | olive baboon | 101008749 | XM_021933040.1 | |
| *Theropithecus gelada* | gelada | 112615413 | XM_025372062.1 | |
| *Chlorocebus sabaeus* | vervet | 103231639 | XM_007991113.1 | |
| *Alouatta palliata* | mantled howler monkey | N/A | N/A | unpublished draft genome, sequence in supplemental file |
| *Aotus nancymaae* | Ma's night monkey | 105705080 | XM_012434682.2 | |
| *Cebus capucinus imitator* | white-faced capuchin | 108291904 | XM_017512376.1 | |
| *Sapajus apella* | tufted capuchin | 116556688 | XM_032285963.1 | |
| *Saimiri boliviensis* | Bolivian squirrel monkey | 101045190 | XM_010336623.1 | |
| *Callithrix jacchus* | common marmoset | 100408882 | XM_017968359.1 | |
| *Carlito syrichta* | Philippine tarsier | 103267011 | XM_008064619.1 | |
| *Microcebus murinus* | gray mouse lemur | 105882317 | XM_020285237.1 | Exon 15 manually corrected, NCBI |

| | | | | |
|---|---|---|---|---|
| | | | | annotation incorrect |
| *Propithecus coquereli* | Coquerel's sifaka | 105805773 | XM_012638731.1 | |
| *Otolemur garnettii* | Northern greater galago | 100951881 | XM_003791864.2 | |
| *Eulemur flavifrons* | blue-eyed black lemur | N/A | LGHW01000591.1, scaffold 590 | *ACE2* not annotated, identified via BLAST |
| *Daubentonia madagascariensis* | aye-aye | N/A | PVJZ01006595.1, scaffold 13170 | *ACE2* not annotated, identified via BLAST |
| *Rhinolophus sinicus* | Chinese rufous horseshoe bat | N/A | GQ999933.1 | From [1] |
| *Rhinolophus pusillus* | least horseshoe bat | N/A | GQ999938.1 | |
| *Rhinolophus macrotis* | big-eared horseshoe bat | N/A | GQ999932.1 | |
| *Rhinolophus pearsonii* | Pearson's horseshoe bat | N/A | EF569964.1 | |
| *Rhinolophus ferrumequinum* | greater horseshoe bat | N/A | GQ999931.1 | |
| *Myotis daubentonii* | Daubenton's bat | N/A | GQ999937.1 | |
| *Hipposideros pratti* | | N/A | GQ999934.1 | |
| *Felis catus* | domestic cat | 554349 | NM_001039456.1 | |
| *Canis lupus* | domestic dog | 480847 | NM_001165260.1 | |
| *Sus scrofa* | domestic pig | 100144303 | NM_001123070.1 | |
| *Mustela putorius* | ferret | 101673097 | NM_001310190.1 | |
| *Manis javanica* | Malayan pangolin | 108390919 | XM_017650257.1 | |

**Supplementary Table S2.** Results of alanine scanning mutagenesis experiments predicting critical binding sites between ACE2 and Sars-CoV-2 receptor binding domain. Residues whose mutation to alanine decrease the binding energy by $\Delta\Delta G_{bind}$ ≥1.0 kcal/mol are considered to be significant for binding (in bold). [+]Denotes sites also implicated by Yan et al.[2]

| Residue site | ΔΔG (kcal/mol)[a] |
|---|---|
| **41[+]** | **4.1** |
| **355** | **3.5** |
| **42[+]** | **2.3** |
| **83** | **2.1** |
| **357[+]** | **2** |
| **38** | **1.4** |
| **37** | **1.2** |
| **24[+]** | **1.1** |
| **353[+]** | **1.1** |
| 27 | 0.7 |
| 34[+] | 0.7 |
| 31 | 0.6 |
| 30[+] | 0.6 |
| 35 | 0.5 |
| 79 | 0.5 |
| 45 | 0.5 |
| 28 | 0.3 |
| 82[+] | 0.2 |
| 330 | 0.2 |
| 351 | 0.03 |

[a]The computational alanine mutagenesis analysis was performed with Rosetta Software and PDB file 6M0J.

**Supplementary Table S3**. Pairwise identities of ACE2 nucleotide sequences between all species in our study. Number indicates the percentage of bases that are identical across the full length of the gene sequence.

| | *Alouatta palliata* | *A. nancymaae* | *C. capucinus* | *S. apella* | *S. boliviensis* | *C. jacchus* | *C. atys* | *M. leucophaeus* | *M. fascicularis* | *M. nemestrina* | *P. anubis* | *T. gelada* | *M. mulatta* | *P. tephrosceles* | *R. roxellana* | *C. sabaeus* | *G. gorilla* | *H. sapiens* | *P. paniscus* | *P. troglodytes* | *P. abelii* | *H. moloch* | *N. leucogenys* | *D. madagascar-iensis* | *E. flavifrons* | *P. coquereli* | *M. murinus* | *O. garnettii* | *C. syrichta* | *C. lupus* | *F. catus* | *M. putorius* | *M. javanica* | *H. pratti* | *R. ferrum-equinum* | *R. macrotis* | *R. pusillus* | *R. sinicus* | *R. pearsonii* | *M. daubentonii* |
|---|---|---|---|---|---|---|---|---|---|---|---|---|---|---|---|---|---|---|---|---|---|---|---|---|---|---|---|---|---|---|---|---|---|---|---|---|---|---|---|---|
| *Aotus nancymaae* | 97.35 | | | | | | | | | | | | | | | | | | | | | | | | | | | | | | | | | | | | | | | |
| *Cebus capucinus* | 97.28 | 97.78 | | | | | | | | | | | | | | | | | | | | | | | | | | | | | | | | | | | | | | |
| *Sapajus apella* | 97.48 | 97.97 | 99.47 | | | | | | | | | | | | | | | | | | | | | | | | | | | | | | | | | | | | | |
| *Saimiri boliviensis* | 97.11 | 97.93 | 98.03 | 98.14 | | | | | | | | | | | | | | | | | | | | | | | | | | | | | | | | | | | | |
| *Callithrix jacchus* | 97.19 | 97.68 | 97.61 | 97.85 | 97.52 | | | | | | | | | | | | | | | | | | | | | | | | | | | | | | | | | | | |
| *Cercocebus atys* | 94.62 | 94.5 | 94.51 | 94.71 | 94.67 | 94.21 | | | | | | | | | | | | | | | | | | | | | | | | | | | | | | | | | | |
| *Mandrillus leucophaeus* | 94.54 | 94.42 | 94.43 | 94.62 | 94.58 | 94.13 | 99.67 | | | | | | | | | | | | | | | | | | | | | | | | | | | | | | | | | |
| *Macaca fascicularis* | 94.79 | 94.67 | 94.55 | 94.75 | 94.75 | 94.29 | 99.42 | 99.34 | | | | | | | | | | | | | | | | | | | | | | | | | | | | | | | | |
| *Macaca nemestrina* | 94.87 | 94.75 | 94.63 | 94.83 | 94.83 | 94.38 | 99.5 | 99.42 | 99.92 | | | | | | | | | | | | | | | | | | | | | | | | | | | | | | | |
| *Papio anubis* | 94.83 | 94.62 | 94.59 | 94.79 | 94.79 | 94.33 | 99.5 | 99.42 | 99.67 | 99.75 | | | | | | | | | | | | | | | | | | | | | | | | | | | | | | |
| *Theropithecus gelada* | 94.71 | 94.58 | 94.55 | 94.75 | 94.83 | 94.29 | 99.59 | 99.5 | 99.5 | 99.59 | 99.75 | | | | | | | | | | | | | | | | | | | | | | | | | | | | | |
| *Macaca mulatta* | 94.67 | 94.54 | 94.43 | 94.62 | 94.54 | 94.17 | 99.17 | 99.09 | 99.59 | 99.67 | 99.42 | 99.26 | | | | | | | | | | | | | | | | | | | | | | | | | | | | |
| *Piliocolobus tephrosceles* | 94.75 | 94.62 | 94.59 | 94.79 | 94.79 | 94.33 | 98.76 | 98.59 | 98.84 | 98.92 | 98.92 | 98.84 | 98.68 | | | | | | | | | | | | | | | | | | | | | | | | | | | |
| *Rhinopithecus roxellana* | 94.79 | 94.54 | 94.59 | 94.79 | 94.79 | 94.33 | 98.92 | 98.76 | 98.97 | 99.05 | 99.05 | 98.97 | 98.72 | 99.05 | | | | | | | | | | | | | | | | | | | | | | | | | | |
| *Chlorocebus sabaeus* | 94.54 | 94.33 | 94.34 | 94.54 | 94.54 | 94.13 | 98.97 | 98.88 | 99.01 | 99.09 | 99.05 | 99.05 | 98.76 | 98.51 | 98.59 | | | | | | | | | | | | | | | | | | | | | | | | | |
| *Gorilla gorilla* | 94.58 | 94.33 | 94.34 | 94.54 | 94.71 | 94.25 | 96.94 | 96.73 | 96.94 | 97.02 | 97.02 | 97.02 | 96.69 | 96.98 | 97.06 | 96.77 | | | | | | | | | | | | | | | | | | | | | | | | |
| *Homo sapiens* | 94.62 | 94.54 | 94.55 | 94.75 | 94.83 | 94.46 | 97.02 | 96.82 | 97.02 | 97.11 | 97.11 | 97.11 | 96.77 | 97.06 | 97.15 | 96.86 | 99.26 | | | | | | | | | | | | | | | | | | | | | | | |
| *Pan paniscus* | 94.54 | 94.46 | 94.47 | 94.67 | 94.75 | 94.38 | 97.02 | 96.82 | 97.02 | 97.02 | 97.02 | 97.02 | 96.69 | 96.98 | 97.06 | 96.77 | 99.17 | 99.42 | | | | | | | | | | | | | | | | | | | | | | |
| *Pan troglodytes* | 94.54 | 94.46 | 94.47 | 94.67 | 94.75 | 94.38 | 97.02 | 96.82 | 96.94 | 97.02 | 97.02 | 97.02 | 96.69 | 96.98 | 97.06 | 96.77 | 99.17 | 99.42 | 99.92 | | | | | | | | | | | | | | | | | | | | | |
| *Pongo abelii* | 94.62 | 94.38 | 94.39 | 94.58 | 94.71 | 94.38 | 97.15 | 96.94 | 97.15 | 97.23 | 97.23 | 97.19 | 96.9 | 97.15 | 97.27 | 96.82 | 98.3 | 98.47 | 98.39 | 98.39 | | | | | | | | | | | | | | | | | | | | |
| *Hylobates moloch* | 94.83 | 94.67 | 94.51 | 94.71 | 94.83 | 94.42 | 96.94 | 96.73 | 96.94 | 97.02 | 97.02 | 96.98 | 96.69 | 96.94 | 97.06 | 96.65 | 98.22 | 98.3 | 98.3 | 98.3 | 98.26 | | | | | | | | | | | | | | | | | | | |
| *Nomascus leucogenys* | 94.87 | 94.62 | 94.47 | 94.67 | 94.87 | 94.46 | 96.94 | 96.73 | 96.94 | 97.02 | 97.02 | 96.98 | 96.69 | 96.94 | 97.06 | 96.61 | 98.26 | 98.26 | 98.26 | 98.26 | 98.35 | 99.26 | | | | | | | | | | | | | | | | | | |
| *Daubentonia madagascariensis* | 89.66 | 89.58 | 89.42 | 89.74 | 89.66 | 89.37 | 90.24 | 90.16 | 90.24 | 90.32 | 90.32 | 90.36 | 90.07 | 90.16 | 90.28 | 90.16 | 89.99 | 90.2 | 90.12 | 90.12 | 90.24 | 90.36 | 90.2 | | | | | | | | | | | | | | | | | |
| *Eulemur flavifrons* | 88.67 | 88.5 | 88.31 | 88.54 | 88.63 | 88.46 | 88.96 | 88.96 | 89.25 | 89.25 | 89.21 | 89.25 | 89 | 89 | 88.96 | 88.96 | 89 | 89.12 | 89.12 | 89.04 | 89.08 | 89.08 | 89.25 | 93.22 | | | | | | | | | | | | | | | | |
| *Propithecus coquereli* | 89.16 | 89.16 | 88.89 | 89.12 | 89.04 | 88.96 | 89.66 | 89.66 | 89.74 | 89.83 | 89.74 | 89.78 | 89.58 | 89.7 | 89.62 | 89.62 | 89.5 | 89.7 | 89.7 | 89.7 | 89.66 | 89.7 | 89.7 | 94.54 | 95.37 | | | | | | | | | | | | | | | |
| *Microcebus murinus* | 88.5 | 88.3 | 88.06 | 88.3 | 88.25 | 88.34 | 88.54 | 88.54 | 88.71 | 88.79 | 88.71 | 88.71 | 88.54 | 88.79 | 88.83 | 88.83 | 88.96 | 88.92 | 88.75 | 88.75 | 88.88 | 88.75 | 88.92 | 93.13 | 94.71 | 95.16 | | | | | | | | | | | | | | |
| *Otolemur garnettii* | 86.35 | 86.19 | 86.07 | 86.39 | 86.48 | 86.1 | 86.35 | 86.27 | 86.31 | 86.39 | 86.31 | 86.44 | 86.15 | 86.56 | 86.56 | 86.48 | 86.52 | 86.48 | 86.39 | 86.39 | 86.35 | 86.44 | 86.48 | 90.24 | 89 | 89.91 | 89.29 | | | | | | | | | | | | | |
| *Carlito syrichta* | 87.84 | 87.39 | 87.56 | 87.76 | 87.84 | 87.68 | 87.88 | 87.88 | 88.01 | 88.09 | 88.09 | 88.13 | 87.88 | 88.13 | 88.05 | 87.92 | 88.17 | 87.92 | 88.05 | 88.05 | 87.97 | 87.97 | 88.09 | 87.76 | 86.64 | 87.26 | 86.97 | 84.66 | | | | | | | | | | | | |
| *Canis lupus* | 86.93 | 86.77 | 86.78 | 86.77 | 87.14 | 86.68 | 87.26 | 87.1 | 87.3 | 87.39 | 87.39 | 87.34 | 87.22 | 87.3 | 87.39 | 87.06 | 87.06 | 87.01 | 87.1 | 87.1 | 87.1 | 87.3 | 87.22 | 88.21 | 86.72 | 87.68 | 86.6 | 85.4 | 85.36 | | | | | | | | | | | |
| *Felis catus* | 86.64 | 86.52 | 86.78 | 86.85 | 86.97 | 86.39 | 86.77 | 86.64 | 86.85 | 86.93 | 86.85 | 86.89 | 86.68 | 86.97 | 86.97 | 86.6 | 86.81 | 86.93 | 87.1 | 87.1 | 86.93 | 87.22 | 87.1 | 87.92 | 86.85 | 87.51 | 86.31 | 85.24 | 85.32 | 92.6 | | | | | | | | | | |
| *Mustela putorius furo* | 85.73 | 85.65 | 85.49 | 85.65 | 85.9 | 85.57 | 85.86 | 85.65 | 85.81 | 85.9 | 85.86 | 85.9 | 85.77 | 85.94 | 85.98 | 85.57 | 85.98 | 85.86 | 85.98 | 85.98 | 85.73 | 85.98 | 86.1 | 86.72 | 85.32 | 86.44 | 85.07 | 84.37 | 84.62 | 92.22 | 91.36 | | | | | | | | | |
| *Manis javanica* | 86.06 | 85.81 | 86.03 | 86.23 | 86.06 | 86.02 | 86.6 | 86.48 | 86.6 | 86.68 | 86.77 | 86.64 | 86.56 | 86.68 | 86.72 | 86.39 | 86.6 | 86.68 | 86.6 | 86.6 | 86.72 | 86.72 | 86.68 | 87.68 | 86.02 | 87.26 | 85.81 | 84.78 | 84.99 | 89.16 | 89.16 | 88.3 | | | | | | | | |
| *Hipposideros pratti* | 85.21 | 84.84 | 85.3 | 85.21 | 84.84 | 85.12 | 85.77 | 85.67 | 85.77 | 85.86 | 85.58 | 85.67 | 85.77 | 85.77 | 85.49 | 85.67 | 85.58 | 85.58 | 85.4 | 85.49 | 85.86 | 85.95 | 85.95 | 85.58 | 85.77 | 85.58 | 84.65 | 82.42 | 82.88 | 87.16 | 86.98 | 85.58 | 86.79 | | | | | | | |
| *Rhinolophus ferrumequinum* | 86.1 | 85.73 | 85.83 | 85.98 | 85.86 | 85.61 | 86.68 | 86.48 | 86.68 | 86.77 | 86.68 | 86.77 | 86.64 | 86.77 | 86.97 | 86.64 | 86.23 | 86.39 | 86.48 | 86.48 | 86.52 | 86.48 | 86.48 | 87.26 | 85.61 | 86.89 | 85.4 | 84.33 | 84.04 | 88.5 | 88.46 | 87.01 | 87.3 | 90.98 | | | | | | |
| *Rhinolophus macrotis* | 85.69 | 85.19 | 85.49 | 85.57 | 85.44 | 85.11 | 85.98 | 85.86 | 85.98 | 86.06 | 85.98 | 86.06 | 85.86 | 86.06 | 86.19 | 85.94 | 85.61 | 85.69 | 85.77 | 85.77 | 85.81 | 85.86 | 85.77 | 87.14 | 85.65 | 86.81 | 85.4 | 84.33 | 83.62 | 87.88 | 88.21 | 86.48 | 87.47 | 90.05 | 95.91 | | | | | |
| *Rhinolophus pusillus* | 85.44 | 84.95 | 85.25 | 85.32 | 85.19 | 84.86 | 85.73 | 85.61 | 85.73 | 85.81 | 85.73 | 85.81 | 85.61 | 85.81 | 85.94 | 85.77 | 85.44 | 85.53 | 85.53 | 85.53 | 85.65 | 85.61 | 85.53 | 86.52 | 85.07 | 86.39 | 84.9 | 83.87 | 83.33 | 87.34 | 87.63 | 86.19 | 86.97 | 89.77 | 95.45 | 98.39 | | | | |
| *Rhinolophus sinicus* | 85.69 | 85.11 | 85.33 | 85.48 | 85.36 | 85.03 | 85.77 | 85.65 | 85.73 | 85.81 | 85.73 | 85.86 | 85.61 | 85.81 | 85.94 | 85.81 | 85.44 | 85.53 | 85.53 | 85.53 | 85.57 | 85.61 | 85.53 | 86.56 | 85.07 | 86.23 | 84.86 | 83.79 | 83.29 | 87.43 | 87.76 | 86.06 | 86.81 | 89.86 | 95.45 | 97.52 | 97.02 | | | |
| *Rhinolophus pearsonii* | 85.9 | 85.32 | 85.54 | 85.69 | 85.57 | 85.4 | 85.98 | 85.81 | 85.94 | 86.02 | 85.94 | 86.02 | 85.81 | 86.02 | 86.15 | 85.9 | 85.65 | 85.65 | 85.65 | 85.65 | 85.77 | 85.9 | 85.81 | 86.85 | 85.4 | 86.56 | 85.19 | 84.08 | 83.71 | 88.05 | 88.17 | 86.56 | 87.06 | 90.05 | 95.66 | 97.02 | 96.36 | 96.94 | | |
| *Myotis daubentonii* | 86.7 | 85.77 | 86.23 | 86.33 | 86.05 | 85.95 | 86.6 | 86.7 | 86.6 | 86.7 | 86.6 | 86.7 | 86.6 | 86.7 | 86.7 | 86.7 | 86.33 | 86.33 | 86.14 | 86.23 | 86.51 | 86.6 | 86.79 | 87.44 | 86.7 | 87.16 | 86.33 | 84.65 | 85.4 | 86.33 | 86.6 | 85.3 | 86.05 | 86.6 | 87.53 | 87.63 | 86.98 | 87.35 | 86.98 | |
| *Sus scrofa* | 84.95 | 84.66 | 84.71 | 84.82 | 84.7 | 84.49 | 84.57 | 84.37 | 84.53 | 84.62 | 84.57 | 84.66 | 84.49 | 84.62 | 84.74 | 84.37 | 84.74 | 84.66 | 84.66 | 84.66 | 84.7 | 84.9 | 84.9 | 85.36 | 84.86 | 85.44 | 84.33 | 82.59 | 82.59 | 86.19 | 87.14 | 86.15 | 85.24 | 84.19 | 85.61 | 85.32 | 84.99 | 84.74 | 84.99 | 83.72 |

**Supplementary Table S4**. Pairwise identities of ACE2 protein sequences between all species in our study. Number indicates the percentage of residues that are identical across the full length of the protein sequence.

| | *Alouatta palliata* | *A. nancymaae* | *C. capucinus* | *S. apella* | *S. boliviensis* | *C. jacchus* | *C. atys* | *M. leucophaeus* | *M. fascicularis* | *M. nemestrina* | *M. mulatta* | *P. anubis* | *T. gelada* | *R. roxellana* | *C. sabaeus* | *P. tephrosceles* | *G. gorilla* | *H. sapiens* | *P. paniscus* | *P. troglodytes* | *H. moloch* | *N. leucogenys* | *P. abelii* | *D. madagascarie nsis* | *E. flavifrons* | *M. murinus* | *O. garnettii* | *P. coquereli* | *C. syrichta* | *C. lupus* | *F. catus* | *M. putorius* | *M. javanica* | *S. scrofa* | *H. pratti* | *R. ferrumequinu m* | *R. macrotis* | *R. pusillus* | *R. sinicus* | *R. pearsonii* |
|---|---|---|---|---|---|---|---|---|---|---|---|---|---|---|---|---|---|---|---|---|---|---|---|---|---|---|---|---|---|---|---|---|---|---|---|---|---|---|---|---|
| *Aotus nancymaae* | 96.89 | | | | | | | | | | | | | | | | | | | | | | | | | | | | | | | | | | | | | | | |
| *Cebus capucinus* | 97.02 | 97.89 | | | | | | | | | | | | | | | | | | | | | | | | | | | | | | | | | | | | | | |
| *Sapajus apella* | 97.02 | 97.89 | 99.51 | | | | | | | | | | | | | | | | | | | | | | | | | | | | | | | | | | | | | |
| *Saimiri boliviensis* | 95.9 | 96.89 | 97.77 | 97.52 | | | | | | | | | | | | | | | | | | | | | | | | | | | | | | | | | | | | |
| *Callithrix jacchus* | 95.78 | 96.77 | 97.15 | 97.14 | 96.27 | | | | | | | | | | | | | | | | | | | | | | | | | | | | | | | | | | | |
| *Cercocebus atys* | 91.55 | 92.05 | 92.06 | 92.05 | 91.68 | 90.68 | | | | | | | | | | | | | | | | | | | | | | | | | | | | | | | | | | |
| *Mandrillus leucophaeus* | 91.3 | 91.8 | 91.81 | 91.8 | 91.43 | 90.43 | 99.63 | | | | | | | | | | | | | | | | | | | | | | | | | | | | | | | | | |
| *Macaca fascicularis* | 91.55 | 92.05 | 92.06 | 92.05 | 91.55 | 90.68 | 99.01 | 98.88 | | | | | | | | | | | | | | | | | | | | | | | | | | | | | | | | |
| *Macaca nemestrina* | 91.68 | 92.17 | 92.18 | 92.17 | 91.68 | 90.81 | 99.13 | 99.01 | 99.88 | | | | | | | | | | | | | | | | | | | | | | | | | | | | | | | |
| *Macaca mulatta* | 91.3 | 91.8 | 91.81 | 91.8 | 91.3 | 90.43 | 98.76 | 98.63 | 99.5 | 99.63 | | | | | | | | | | | | | | | | | | | | | | | | | | | | | | |
| *Papio anubis* | 91.55 | 92.05 | 92.06 | 92.05 | 91.55 | 90.68 | 99.01 | 99.01 | 99.25 | 99.38 | 99.01 | | | | | | | | | | | | | | | | | | | | | | | | | | | | | |
| *Theropithecus gelada* | 91.68 | 92.17 | 92.18 | 92.17 | 91.8 | 90.81 | 99.25 | 99.25 | 99.13 | 99.25 | 98.88 | 99.63 | | | | | | | | | | | | | | | | | | | | | | | | | | | | |
| *Rhinopithecus roxellana* | 91.06 | 91.3 | 91.56 | 91.55 | 91.18 | 90.19 | 98.14 | 98.01 | 98.26 | 98.39 | 98.01 | 98.26 | 98.39 | | | | | | | | | | | | | | | | | | | | | | | | | | | |
| *Chlorocebus sabaeus* | 91.3 | 91.55 | 91.56 | 91.55 | 91.18 | 90.56 | 98.26 | 98.14 | 98.39 | 98.51 | 98.14 | 98.26 | 98.39 | 97.39 | | | | | | | | | | | | | | | | | | | | | | | | | | |
| *Piliocolobus tephrosceles* | 91.3 | 91.55 | 91.81 | 91.8 | 91.55 | 90.43 | 97.76 | 97.64 | 97.89 | 98.01 | 97.64 | 97.89 | 98.01 | 97.89 | 97.14 | | | | | | | | | | | | | | | | | | | | | | | | | |
| *Gorilla gorilla* | 92.42 | 91.93 | 92.3 | 92.3 | 91.8 | 91.43 | 95.28 | 95.03 | 95.28 | 95.4 | 95.03 | 95.28 | 95.4 | 95.4 | 94.91 | 95.16 | | | | | | | | | | | | | | | | | | | | | | | | |
| *Homo sapiens* | 92.17 | 92.17 | 92.55 | 92.55 | 92.05 | 91.68 | 95.16 | 94.91 | 95.16 | 95.28 | 94.91 | 95.28 | 95.28 | 95.28 | 94.78 | 95.03 | 99.01 | | | | | | | | | | | | | | | | | | | | | | | |
| *Pan paniscus* | 92.05 | 91.93 | 92.3 | 92.3 | 91.8 | 91.55 | 95.03 | 94.78 | 95.28 | 95.16 | 94.78 | 95.03 | 95.16 | 95.16 | 94.66 | 94.91 | 98.76 | 99.01 | | | | | | | | | | | | | | | | | | | | | | |
| *Pan troglodytes* | 92.05 | 91.93 | 92.3 | 92.3 | 91.8 | 91.55 | 95.03 | 94.78 | 95.03 | 95.16 | 94.78 | 95.03 | 95.16 | 95.16 | 94.66 | 94.91 | 98.76 | 99.01 | 99.75 | | | | | | | | | | | | | | | | | | | | | |
| *Hylobates moloch* | 92.42 | 92.17 | 92.3 | 92.3 | 91.68 | 91.43 | 94.78 | 94.53 | 94.78 | 94.91 | 94.53 | 94.78 | 94.91 | 94.91 | 94.29 | 94.66 | 98.01 | 98.01 | 98.01 | 98.01 | | | | | | | | | | | | | | | | | | | | |
| *Nomascus leucogenys* | 92.3 | 91.93 | 92.06 | 92.05 | 91.68 | 91.18 | 94.66 | 94.41 | 94.53 | 94.66 | 94.29 | 94.53 | 94.78 | 94.78 | 94.16 | 94.53 | 98.01 | 97.76 | 97.76 | 97.76 | 98.76 | | | | | | | | | | | | | | | | | | | |
| *Pongo abelii* | 91.8 | 91.68 | 92.06 | 92.05 | 91.68 | 91.18 | 95.28 | 95.03 | 95.16 | 95.28 | 94.91 | 95.16 | 95.4 | 95.4 | 94.78 | 95.16 | 98.14 | 98.14 | 97.89 | 97.89 | 97.52 | 97.64 | | | | | | | | | | | | | | | | | | |
| *Daubentonia madagascariensis* | 85.84 | 86.09 | 86.47 | 86.34 | 85.71 | 85.34 | 86.83 | 86.71 | 86.71 | 86.83 | 86.46 | 86.83 | 86.96 | 86.46 | 86.21 | 86.34 | 86.34 | 86.83 | 86.58 | 86.58 | 86.58 | 86.21 | 86.34 | | | | | | | | | | | | | | | | | |
| *Eulemur flavifrons* | 83.23 | 83.11 | 83.61 | 83.48 | 82.98 | 82.73 | 84.35 | 84.47 | 84.6 | 84.47 | 84.1 | 84.6 | 84.6 | 83.73 | 84.1 | 84.35 | 84.22 | 84.35 | 84.35 | 84.1 | 84.22 | 84.1 | 83.98 | 87.58 | | | | | | | | | | | | | | | | |
| *Microcebus murinus* | 83.85 | 83.11 | 83.86 | 83.85 | 83.23 | 83.35 | 83.73 | 83.73 | 83.85 | 83.98 | 83.6 | 83.73 | 83.85 | 83.73 | 84.1 | 84.1 | 84.6 | 84.1 | 83.73 | 83.73 | 83.85 | 83.85 | 83.98 | 87.95 | 90.81 | | | | | | | | | | | | | | | |
| *Otolemur garnettii* | 81.74 | 81.74 | 81.99 | 82.11 | 81.49 | 81.12 | 81.61 | 81.49 | 81.74 | 81.86 | 81.49 | 81.74 | 81.86 | 81.74 | 81.49 | 82.36 | 81.74 | 81.86 | 81.74 | 81.74 | 81.61 | 81.37 | 81.49 | 84.6 | 83.11 | 83.85 | | | | | | | | | | | | | | |
| *Propithecus coquereli* | 84.84 | 85.09 | 85.22 | 85.22 | 84.84 | 84.35 | 85.96 | 85.96 | 86.09 | 86.21 | 85.84 | 85.96 | 86.09 | 85.47 | 85.47 | 86.09 | 85.47 | 85.71 | 85.84 | 85.84 | 85.96 | 85.47 | 85.22 | 90.19 | 91.93 | 91.68 | 84.97 | | | | | | | | | | | | | |
| *Carlito syrichta* | 83.73 | 83.35 | 83.61 | 83.73 | 83.48 | 83.35 | 83.85 | 83.73 | 83.85 | 83.98 | 83.6 | 83.85 | 84.1 | 83.73 | 83.73 | 84.35 | 84.72 | 84.1 | 84.22 | 84.22 | 84.1 | 84.35 | 84.22 | 82.36 | 80.99 | 81.74 | 79.88 | 82.11 | | | | | | | | | | | | |
| *Canis lupus* | 81.99 | 82.11 | 82.37 | 82.24 | 81.99 | 81.12 | 83.48 | 83.23 | 83.35 | 83.48 | 83.11 | 83.73 | 83.6 | 83.23 | 83.11 | 83.35 | 83.23 | 83.35 | 83.35 | 83.35 | 83.35 | 82.86 | 83.35 | 82.36 | 80.25 | 81.37 | 80.62 | 82.48 | 79.88 | | | | | | | | | | | |
| *Felis catus* | 83.73 | 84.22 | 84.73 | 84.6 | 84.35 | 83.35 | 84.84 | 84.6 | 84.72 | 84.84 | 84.47 | 84.84 | 84.97 | 84.72 | 84.35 | 84.84 | 84.97 | 85.22 | 85.34 | 85.34 | 85.59 | 85.22 | 84.97 | 84.72 | 81.86 | 82.48 | 81.49 | 84.35 | 80.75 | 90.56 | | | | | | | | | | |
| *Mustela putorius furo* | 81.61 | 82.24 | 82.37 | 82.24 | 82.11 | 81.12 | 82.61 | 82.36 | 82.48 | 82.61 | 82.36 | 82.61 | 82.73 | 82.61 | 82.11 | 82.73 | 82.86 | 82.61 | 82.98 | 82.98 | 82.73 | 82.48 | 82.24 | 82.36 | 79.5 | 80 | 80.25 | 82.61 | 80.62 | 89.57 | 89.69 | | | | | | | | | |
| *Manis javanica* | 83.6 | 83.98 | 84.11 | 84.1 | 83.73 | 82.98 | 85.22 | 84.97 | 85.22 | 85.34 | 84.97 | 85.47 | 85.47 | 85.22 | 84.84 | 85.47 | 84.97 | 84.84 | 84.97 | 84.97 | 84.72 | 84.6 | 84.72 | 84.6 | 81.49 | 82.24 | 81.24 | 83.98 | 81.86 | 86.21 | 87.33 | 86.58 | | | | | | | | |
| *Sus scrofa* | 81.24 | 81.12 | 81.25 | 81.24 | 80.62 | 80.37 | 81.24 | 80.99 | 81.12 | 81.24 | 80.87 | 81.24 | 81.37 | 81.24 | 80.87 | 80.87 | 81.61 | 81.37 | 81.37 | 81.37 | 81.49 | 81.49 | 81.61 | 81.74 | 80.62 | 81.24 | 77.89 | 81.86 | 79.25 | 81.99 | 83.98 | 83.85 | 82.61 | | | | | | | |
| *Hipposideros pratti* | 77.65 | 77.65 | 78.49 | 78.49 | 78.21 | 76.82 | 78.21 | 78.21 | 77.93 | 78.21 | 77.65 | 77.93 | 78.21 | 77.37 | 78.21 | 77.65 | 77.93 | 77.37 | 77.37 | 77.65 | 77.65 | 77.65 | 77.37 | 77.93 | 77.37 | 76.82 | 73.74 | 77.93 | 74.3 | 78.49 | 80.73 | 78.21 | 80.73 | 76.54 | | | | | | |
| *Rhinolophus ferrumequinum* | 80 | 80.37 | 80.75 | 80.75 | 80.37 | 79.75 | 81.74 | 81.49 | 81.61 | 81.74 | 81.37 | 81.74 | 81.86 | 81.99 | 81.49 | 81.49 | 81.24 | 81.49 | 81.61 | 81.61 | 81.37 | 80.99 | 81.49 | 80.12 | 77.27 | 77.14 | 77.64 | 79.25 | 76.77 | 81.74 | 83.73 | 80.99 | 83.11 | 80.37 | 83.52 | | | | | |
| *Rhinolophus macrotis* | 79.88 | 80.12 | 80.75 | 80.75 | 80.37 | 79.5 | 80.87 | 80.62 | 80.75 | 80.87 | 80.5 | 80.87 | 80.99 | 80.87 | 80.62 | 80.62 | 80.62 | 80.87 | 80.99 | 80.99 | 80.75 | 80.37 | 80.75 | 80.5 | 77.76 | 78.51 | 78.26 | 79.88 | 76.02 | 81.74 | 84.35 | 81.12 | 83.73 | 80.5 | 84.36 | 92.42 | | | | |
| *Rhinolophus pusillus* | 79.88 | 79.88 | 80.5 | 80.5 | 79.88 | 79.25 | 80.37 | 80.12 | 80.25 | 80.37 | 80 | 80.37 | 80.5 | 80.37 | 80.37 | 80.12 | 80.37 | 80.5 | 80.62 | 80.62 | 80.5 | 80.25 | 80.37 | 79.75 | 77.39 | 78.14 | 77.52 | 79.25 | 76.27 | 81.61 | 84.1 | 81.49 | 83.35 | 80.99 | 84.08 | 91.8 | 96.77 | | | |
| *Rhinolophus sinicus* | 79.88 | 80 | 80.63 | 80.62 | 80 | 79.38 | 80.25 | 80 | 80.12 | 80.25 | 79.88 | 80.25 | 80.37 | 80.25 | 80.37 | 80 | 80 | 80.12 | 80.25 | 80.25 | 80 | 79.75 | 80 | 79.5 | 77.14 | 77.39 | 77.14 | 78.88 | 76.02 | 80.75 | 83.35 | 80.25 | 82.24 | 79.63 | 84.08 | 92.3 | 95.28 | 94.29 | | |
| *Rhinolophus pearsonii* | 80.87 | 80.87 | 81.5 | 81.49 | 81.12 | 80.37 | 81.12 | 80.87 | 80.99 | 81.12 | 80.75 | 81.12 | 81.24 | 81.12 | 80.87 | 80.87 | 81.37 | 81.37 | 81.49 | 81.49 | 81.49 | 81.12 | 81.24 | 80.37 | 77.76 | 78.39 | 78.26 | 79.88 | 77.27 | 82.11 | 84.22 | 81.24 | 82.98 | 80.37 | 83.8 | 92.3 | 94.91 | 93.42 | 94.78 | |
| *Myotis daubentonii* | 79.89 | 79.89 | 80.17 | 80.17 | 79.89 | 78.77 | 80.17 | 80.17 | 79.89 | 80.17 | 79.61 | 79.89 | 80.17 | 79.33 | 79.89 | 79.61 | 79.33 | 78.77 | 78.77 | 79.05 | 79.05 | 79.05 | 78.77 | 80.17 | 79.33 | 78.49 | 77.09 | 79.61 | 77.09 | 77.37 | 80.45 | 78.77 | 79.05 | 77.09 | 79.05 | 77.93 | 79.89 | 79.05 | 78.77 | 78.21 |

**Supplementary Table S5.** Pairwise similarities of ACE2 protein sequences between all species in our study. Number indicates the percentage of residues that are similar (score ≥1 in the BLOSUM62 matrix) across the full length of the protein sequence.

| | Alouatta palliata | A. nancymaae | C. capucinus | S. apella | S. boliviensis | C. jacchus | C. atys | M. leucophaeus | M. fascicularis | M. nemestrina | M. mulatta | P. anubis | T. gelada | R. roxellana | C. sabaeus | P. tephrosceles | G. gorilla | H. sapiens | P. paniscus | P. troglodytes | H. moloch | N. leucogenys | P. abelii | D. madagascar-iensis | E. flavifrons | M. murinus | O. garnettii | P. coquereli | C. syrichta | C. lupus | F. catus | M. putorius | M. javanica | S. scrofa | H. pratti | R. ferrum-equinum | R. macrotis | R. pusillus | R. sinicus | R. pearsonii |
|---|---|---|---|---|---|---|---|---|---|---|---|---|---|---|---|---|---|---|---|---|---|---|---|---|---|---|---|---|---|---|---|---|---|---|---|---|---|---|---|---|
| Aotus nancymaae | 98.26 | | | | | | | | | | | | | | | | | | | | | | | | | | | | | | | | | | | | | | | |
| Cebus capucinus | 98.63 | 98.63 | | | | | | | | | | | | | | | | | | | | | | | | | | | | | | | | | | | | | | |
| Sapajus apella | 98.76 | 98.76 | 99.88 | | | | | | | | | | | | | | | | | | | | | | | | | | | | | | | | | | | | | |
| Saimiri boliviensis | 97.76 | 98.01 | 98.39 | 98.51 | | | | | | | | | | | | | | | | | | | | | | | | | | | | | | | | | | | | |
| Callithrix jacchus | 97.76 | 97.89 | 98.26 | 98.39 | 97.76 | | | | | | | | | | | | | | | | | | | | | | | | | | | | | | | | | | | |
| Cercocebus atys | 96.27 | 96.15 | 96.15 | 96.27 | 95.78 | 95.4 | | | | | | | | | | | | | | | | | | | | | | | | | | | | | | | | | | |
| Mandrillus leucophaeus | 96.02 | 95.9 | 95.9 | 96.02 | 95.65 | 95.16 | 99.63 | | | | | | | | | | | | | | | | | | | | | | | | | | | | | | | | | |
| Macaca fascicularis | 96.52 | 96.4 | 96.15 | 96.27 | 96.02 | 95.53 | 99.38 | 99.25 | | | | | | | | | | | | | | | | | | | | | | | | | | | | | | | | |
| Macaca nemestrina | 96.52 | 96.4 | 96.15 | 96.27 | 96.02 | 95.53 | 99.38 | 99.25 | 100 | | | | | | | | | | | | | | | | | | | | | | | | | | | | | | | |
| Macaca mulatta | 96.27 | 96.15 | 95.9 | 96.02 | 95.78 | 95.28 | 99.13 | 99.01 | 99.75 | 99.75 | | | | | | | | | | | | | | | | | | | | | | | | | | | | | | |
| Papio anubis | 96.65 | 96.4 | 96.27 | 96.4 | 96.15 | 95.65 | 99.38 | 99.25 | 99.75 | 99.75 | 99.5 | | | | | | | | | | | | | | | | | | | | | | | | | | | | | |
| Theropithecus gelada | 96.65 | 96.52 | 96.4 | 96.52 | 96.27 | 95.78 | 99.5 | 99.38 | 99.63 | 99.63 | 99.38 | 99.88 | | | | | | | | | | | | | | | | | | | | | | | | | | | | |
| Rhinopithecus roxellana | 96.4 | 96.02 | 96.15 | 96.27 | 96.02 | 95.53 | 98.63 | 98.51 | 99.01 | 99.01 | 98.76 | 99.01 | 99.13 | | | | | | | | | | | | | | | | | | | | | | | | | | | |
| Chlorocebus sabaeus | 96.02 | 95.65 | 95.65 | 95.78 | 95.4 | 94.91 | 98.88 | 98.76 | 99.01 | 99.01 | 98.76 | 98.76 | 98.88 | 98.26 | | | | | | | | | | | | | | | | | | | | | | | | | | |
| Piliocolobus tephrosceles | 96.27 | 95.9 | 96.02 | 96.15 | 95.9 | 95.4 | 98.51 | 98.39 | 98.88 | 98.88 | 98.63 | 98.88 | 99.01 | 99.13 | 98.14 | | | | | | | | | | | | | | | | | | | | | | | | | |
| Gorilla gorilla | 96.27 | 95.53 | 95.9 | 96.02 | 95.65 | 95.4 | 97.52 | 97.27 | 97.76 | 97.76 | 97.52 | 97.76 | 97.89 | 98.01 | 97.14 | 97.89 | | | | | | | | | | | | | | | | | | | | | | | | |
| Homo sapiens | 96.02 | 95.53 | 95.9 | 96.02 | 95.65 | 95.4 | 97.52 | 97.27 | 97.76 | 97.76 | 97.52 | 97.64 | 97.89 | 98.01 | 97.14 | 97.89 | 99.5 | | | | | | | | | | | | | | | | | | | | | | | |
| Pan paniscus | 95.9 | 95.4 | 95.78 | 95.9 | 95.53 | 95.4 | 97.52 | 97.27 | 97.64 | 97.64 | 97.39 | 97.64 | 97.76 | 97.89 | 97.14 | 97.76 | 99.38 | 99.38 | | | | | | | | | | | | | | | | | | | | | | |
| Pan troglodytes | 95.9 | 95.4 | 95.78 | 95.9 | 95.53 | 95.4 | 97.52 | 97.27 | 97.64 | 97.64 | 97.39 | 97.64 | 97.76 | 97.89 | 97.14 | 97.76 | 99.38 | 99.38 | 100 | | | | | | | | | | | | | | | | | | | | | |
| Hylobates moloch | 96.15 | 95.65 | 96.02 | 96.15 | 95.78 | 95.53 | 97.64 | 97.39 | 97.89 | 97.89 | 97.64 | 97.89 | 98.01 | 98.14 | 97.27 | 98.01 | 99.13 | 99.13 | 99.01 | 99.01 | | | | | | | | | | | | | | | | | | | | |
| Nomascus leucogenys | 96.27 | 95.78 | 96.15 | 96.27 | 95.9 | 95.65 | 97.76 | 97.52 | 98.01 | 98.01 | 97.76 | 98.01 | 98.14 | 98.26 | 97.39 | 98.14 | 99.25 | 99.25 | 99.13 | 99.13 | 99.75 | | | | | | | | | | | | | | | | | | | |
| Pongo abelii | 96.27 | 95.78 | 96.15 | 96.27 | 95.9 | 95.65 | 97.76 | 97.52 | 98.01 | 98.01 | 97.76 | 98.01 | 98.14 | 98.26 | 97.39 | 98.14 | 99.25 | 99.25 | 99.13 | 99.13 | 99.38 | 99.5 | | | | | | | | | | | | | | | | | | |
| Daubentonia madagascarien | 92.8 | 92.55 | 92.67 | 92.8 | 92.17 | 92.05 | 93.29 | 93.17 | 93.54 | 93.54 | 93.29 | 93.66 | 93.79 | 93.42 | 92.8 | 93.29 | 93.04 | 93.29 | 92.92 | 92.92 | 93.17 | 93.17 | 93.29 | | | | | | | | | | | | | | | | | |
| Eulemur flavifrons | 91.8 | 91.93 | 91.93 | 92.05 | 91.68 | 91.3 | 92.55 | 92.42 | 93.04 | 93.04 | 92.8 | 92.92 | 92.8 | 92.3 | 92.17 | 92.42 | 91.8 | 92.05 | 91.68 | 91.68 | 92.17 | 92.17 | 92.05 | 94.16 | | | | | | | | | | | | | | | | |
| Microcebus murinus | 91.3 | 91.18 | 91.3 | 91.43 | 91.06 | 90.68 | 91.8 | 91.68 | 92.05 | 92.05 | 91.8 | 92.17 | 92.3 | 92.3 | 92.05 | 92.67 | 92.05 | 92.05 | 91.68 | 91.68 | 91.8 | 91.93 | 92.05 | 94.04 | 95.16 | | | | | | | | | | | | | | | |
| Otolemur garnettii | 89.69 | 89.32 | 89.32 | 89.44 | 89.19 | 88.82 | 89.19 | 89.07 | 89.32 | 89.32 | 89.07 | 89.44 | 89.57 | 89.57 | 88.94 | 89.94 | 89.44 | 89.44 | 89.07 | 89.07 | 89.32 | 89.32 | 89.44 | 91.8 | 90.81 | 91.68 | | | | | | | | | | | | | | |
| Propithecus coquereli | 91.93 | 92.3 | 91.8 | 91.93 | 91.8 | 91.43 | 92.92 | 92.8 | 93.17 | 93.17 | 92.92 | 93.29 | 93.42 | 92.8 | 92.55 | 93.29 | 92.55 | 92.8 | 92.67 | 92.67 | 92.8 | 92.8 | 92.8 | 94.91 | 96.27 | 96.15 | 91.55 | | | | | | | | | | | | | |
| Carlito syrichta | 91.43 | 90.93 | 91.43 | 91.55 | 91.3 | 91.18 | 91.18 | 91.06 | 91.43 | 91.43 | 91.18 | 91.55 | 91.68 | 91.68 | 90.93 | 92.05 | 92.05 | 91.8 | 91.93 | 91.93 | 91.8 | 91.8 | 91.68 | 90.56 | 89.19 | 89.81 | 88.57 | 89.94 | | | | | | | | | | | | |
| Canis lupus | 91.3 | 91.06 | 91.18 | 91.3 | 90.93 | 90.56 | 92.05 | 91.8 | 92.17 | 92.17 | 91.93 | 92.3 | 92.42 | 92.3 | 91.55 | 92.3 | 91.8 | 91.68 | 91.68 | 91.68 | 92.05 | 91.93 | 92.05 | 91.43 | 89.94 | 90.43 | 89.57 | 91.06 | 89.19 | | | | | | | | | | | |
| Felis catus | 92.42 | 92.17 | 92.3 | 92.42 | 92.17 | 91.55 | 92.55 | 92.3 | 92.67 | 92.67 | 92.42 | 92.8 | 92.92 | 92.92 | 91.93 | 92.92 | 92.3 | 92.3 | 92.17 | 92.17 | 92.55 | 92.42 | 92.55 | 92.67 | 91.3 | 91.43 | 90.56 | 92.42 | 89.57 | 94.53 | | | | | | | | | | |
| Mustela putorius furo | 91.55 | 91.18 | 91.18 | 91.3 | 91.18 | 90.68 | 91.8 | 91.55 | 91.8 | 91.8 | 91.43 | 91.93 | 92.05 | 91.93 | 91.3 | 92.05 | 91.8 | 91.55 | 91.68 | 91.68 | 91.68 | 91.55 | 91.55 | 91.43 | 90.43 | 89.81 | 89.19 | 91.43 | 89.32 | 94.16 | 95.03 | | | | | | | | | |
| Manis javanica | 91.3 | 91.06 | 91.06 | 91.18 | 90.81 | 90.43 | 91.55 | 91.3 | 91.68 | 91.68 | 91.43 | 91.8 | 91.93 | 91.93 | 91.18 | 92.05 | 91.55 | 91.3 | 91.18 | 91.18 | 91.43 | 91.55 | 91.55 | 92.3 | 90.06 | 90.93 | 89.69 | 91.06 | 88.94 | 91.93 | 92.42 | 93.04 | | | | | | | | |
| Sus scrofa | 89.94 | 89.81 | 90.06 | 90.19 | 89.57 | 89.57 | 90.68 | 90.43 | 90.81 | 90.81 | 90.56 | 90.93 | 90.93 | 90.93 | 90.43 | 90.56 | 90.68 | 90.68 | 90.56 | 90.56 | 90.81 | 90.93 | 90.81 | 90.56 | 90.19 | 90.31 | 88.32 | 90.31 | 87.83 | 90.56 | 91.93 | 90.81 | 90.43 | | | | | | | |
| Hipposideros pratti | 86.31 | 85.75 | 86.31 | 86.31 | 86.03 | 85.47 | 86.87 | 86.87 | 86.87 | 86.87 | 86.31 | 86.87 | 86.87 | 86.31 | 86.59 | 86.03 | 86.03 | 85.75 | 85.75 | 85.75 | 86.31 | 86.31 | 85.75 | 87.15 | 86.87 | 86.03 | 84.36 | 86.31 | 82.96 | 87.43 | 87.99 | 86.59 | 87.71 | 85.75 | | | | | | |
| Rhinolophus ferrumequinum | 90.19 | 90.06 | 90.19 | 90.31 | 89.94 | 89.69 | 90.81 | 90.56 | 90.81 | 90.81 | 90.56 | 90.93 | 91.06 | 90.93 | 90.56 | 90.81 | 90.31 | 90.31 | 90.19 | 90.19 | 90.31 | 90.43 | 90.43 | 89.57 | 88.7 | 88.57 | 88.32 | 89.19 | 87.08 | 89.81 | 91.18 | 89.32 | 89.94 | 89.32 | 92.18 | | | | | |
| Rhinolophus macrotis | 89.94 | 89.69 | 90.06 | 90.19 | 89.81 | 89.32 | 90.56 | 90.31 | 90.56 | 90.56 | 90.31 | 90.68 | 90.81 | 90.68 | 90.31 | 90.56 | 90.06 | 90.06 | 89.94 | 89.94 | 90.06 | 90.19 | 90.06 | 89.69 | 88.7 | 89.07 | 88.82 | 89.44 | 86.58 | 89.94 | 91.06 | 89.57 | 90.43 | 88.7 | 91.34 | 95.65 | | | | |
| Rhinolophus pusillus | 90.19 | 89.94 | 90.31 | 90.43 | 89.81 | 89.57 | 90.56 | 90.31 | 90.56 | 90.56 | 90.31 | 90.68 | 90.81 | 90.68 | 90.56 | 90.56 | 89.94 | 90.06 | 89.94 | 89.94 | 89.94 | 90.06 | 90.06 | 89.44 | 88.45 | 88.82 | 88.32 | 88.94 | 86.83 | 89.81 | 90.81 | 89.81 | 90.19 | 88.82 | 91.06 | 95.53 | 97.89 | | | |
| Rhinolophus sinicus | 90.43 | 90.19 | 90.56 | 90.68 | 89.94 | 89.81 | 90.19 | 89.94 | 90.19 | 90.19 | 89.94 | 90.31 | 90.43 | 90.31 | 90.19 | 90.19 | 89.69 | 89.69 | 89.57 | 89.57 | 89.69 | 89.81 | 89.69 | 89.07 | 88.2 | 88.32 | 87.45 | 88.82 | 86.34 | 89.19 | 90.43 | 88.94 | 89.19 | 88.57 | 89.11 | 95.03 | 97.39 | 97.02 | | |
| Rhinolophus pearsonii | 90.56 | 90.31 | 90.68 | 90.81 | 90.43 | 90.19 | 91.06 | 90.81 | 91.06 | 91.06 | 90.81 | 91.18 | 91.3 | 91.18 | 90.81 | 91.06 | 90.56 | 90.56 | 90.43 | 90.43 | 90.56 | 90.68 | 90.56 | 89.69 | 88.45 | 88.82 | 88.7 | 89.44 | 86.96 | 90.81 | 91.3 | 89.69 | 89.69 | 88.82 | 89.94 | 95.53 | 97.39 | 96.27 | 97.14 | |
| Myotis daubentonii | 87.43 | 86.87 | 86.87 | 87.15 | 86.59 | 85.75 | 87.71 | 87.71 | 87.71 | 87.71 | 87.15 | 87.71 | 87.71 | 87.15 | 87.43 | 86.87 | 86.31 | 86.03 | 86.03 | 86.03 | 86.31 | 86.31 | 86.03 | 88.55 | 88.83 | 87.71 | 86.59 | 88.27 | 85.75 | 87.15 | 88.27 | 87.99 | 87.71 | 87.71 | 87.15 | 88.27 | 87.99 | 88.55 | 87.71 | 86.87 |

**Supplementary Table S6.** Full results of the codeml analyses of adaptive evolution across *ACE2* gene sequences.

| codeml analysis | Foreground branch | Model | kappa | treeLength | number of parameters | omega | proportion | lnL | LRT | p | positively selected sites |
|---|---|---|---|---|---|---|---|---|---|---|---|
| branch-site model | platyrrhine | null | 3.08706 | 3.33363 | 79 | background: $\omega_0$ = 0.07549, $\omega_1$ = 1.00000, $\omega_{2a}$ = 0.07549, $\omega_{2b}$ = 1.00000; foreground: $\omega_0$ = 0.07549, $\omega_1$ = 1.00000, $\omega_{2a}$ = 1.00000, $\omega_{2b}$ = 1.00000 | p0 = 0.63315, p1 = 0.35752, p2a = 0.00596, p2b = 0.00337 | -15583.14271 | | | |
| branch-site model | platyrrhine | alternative | 3.08849 | 3.3349 | 80 | background: $\omega_0$ = 0.07584, $\omega_1$ = 1.00000, $\omega_{2a}$ = 0.07584, $\omega_{2b}$ = 1.00000; foreground: $\omega_0$ = 0.07584, $\omega_1$ = 1.00000, $\omega_{2a}$ = 6.21808, $\omega_{2b}$ = 6.21808 | p0 = 0.63821, p1 = 0.35912, p2a = 0.00171, p2b = 0.00096 | -15582.82647 | 0.632472 | 0.4264500154 | n/a |
| branch-site model | bats | null | 3.08122 | 3.33427 | 79 | background: $\omega_0$ = 0.07232, $\omega_1$ = 1.00000, $\omega_{2a}$ = 0.07232, $\omega_{2b}$ = 1.00000; foreground: $\omega_0$ = 0.07232, $\omega_1$ = 1.00000, $\omega_{2a}$ = 1.00000, $\omega_{2b}$ = 1.00000 | p0 = 0.58383, p1 = 0.32072, p2a = 0.06161, p2b = 0.03384 | -15579.5976 | | | |
| branch-site model | bats | alternative | 3.12988 | 3.35579 | 80 | background: $\omega_0$ = 0.07530, $\omega_1$ = 1.00000, $\omega_{2a}$ = 0.07530, $\omega_{2b}$ = 1.00000; foreground: $\omega_0$ = 0.07530, $\omega_1$ = 1.00000, $\omega_{2a}$ = 10.53445, $\omega_{2b}$ = 10.53445 | p0 = 0.62612, p1 = 0.33778, p2a = 0.02345, p2b = 0.01265 | -15558.2729 | 42.649394 | 0 | 7L (0.917), **24Q (0.998**)**, 27T (0.718), **31E (0.959*)**, 34H (0.887), **35E (0.974*)**, 42E (0.652), 91L (0.888), **298V (0.959*)**, 478W (0.505), 483E (0.768), 549E (0.807), 565P (0.939), **568L (0.998**)**, 569A (0.764), **575G (0.965*)**, 658V (0.810), 771K (0.570) |
| branch-site model | catarrhines | null | 3.08696 | 3.33682 | 79 | background: $\omega_0$ = 0.07190, $\omega_1$ = 1.00000, $\omega_{2a}$ = 0.07190, $\omega_{2b}$ = 1.00000; foreground: $\omega_0$ = 0.07190, $\omega_1$ = 1.00000, $\omega_{2a}$ = 1.00000, $\omega_{2b}$ = 1.00000 | p0 = 0.59837, p1 = 0.33784, p2a = 0.04077, p2b = 0.02302 | -15581.0318 | | | |
| branch-site model | catarrhines | alternative | 3.10444 | 3.33652 | 80 | background: $\omega_0$ = 0.07498, $\omega_1$ = 1.00000, $\omega_{2a}$ = 0.07498, $\omega_{2b}$ = 1.00000; foreground: $\omega_0$ = 0.07498, $\omega_1$ = 1.00000, $\omega_{2a}$ = 8.98776, $\omega_{2b}$ = 8.98776 | p0 = 0.63077, p1 = 0.35602, p2a = 0.00845, p2b = 0.00477 | -15573.75876 | 14.546074 | 0.000136773355 | 206D (0.786), **249M (0.962*)**, 338D (0.581), **653A (0.958*)**, 657K (0.511), **658V (0.957*)**, 706M (0.827), 729P (0.798), 732G (0.616), 788K (0.581) |
| branch-site model | strepsirrhines | null | 3.0744 | 3.34127 | 79 | background: $\omega_0$ = 0.06994, $\omega_1$ = 1.00000, $\omega_{2a}$ = 0.06994, $\omega_{2b}$ = 1.00000; foreground: $\omega_0$ = 0.06994, $\omega_1$ = 1.00000, $\omega_{2a}$ = 1.00000, $\omega_{2b}$ = 1.00000 | p0 = 0.59047, p1 = 0.30920, p2a = 0.06585, p2b = 0.03448 | -15576.45584 | | | |
| branch-site model | strepsirrhines | alternative | 3.08067 | 3.34233 | 80 | background: $\omega_0$ = 0.07170, $\omega_1$ = 1.00000, $\omega_{2a}$ = 0.07170, $\omega_{2b}$ = 1.00000; foreground: $\omega_0$ = 0.07170, $\omega_1$ = 1.00000, $\omega_{2a}$ = 1.38404, $\omega_{2b}$ = 1.38404 | p0 = 0.60671, p1 = 0.31603, p2a = 0.05080, p2b = 0.02646 | -15576.03928 | 0.83312 | 0.3613718973 | n/a |
| cladeC model | n/a | M2a_rel (cla | 2.99711 | 3.34639 | 80 | $\omega_0$ = 0.03668, $\omega_1$ = 1.00000, $\omega_2$ = 0.36059 | p0 = 0.50433, p1 = 0.27145, p2 = 0.22423 | -15575.33101 | | | |
| cladeC model | n/a | cladeC | 3.07137 | 3.34517 | 83 | $\omega_0$ = 0.05915, $\omega_1$ = 1.00000, $\omega_2$ = 0.08051, $\omega_3$ = 1.12260, $\omega_4$ = 0.23634, $\omega_5$ = 1.34632 | p0 = 0.58058, p1 = 0.33089, p2-5 = 0.08853 | -15561.96782 | 26.72638 | 0.000006718510 | n/a |

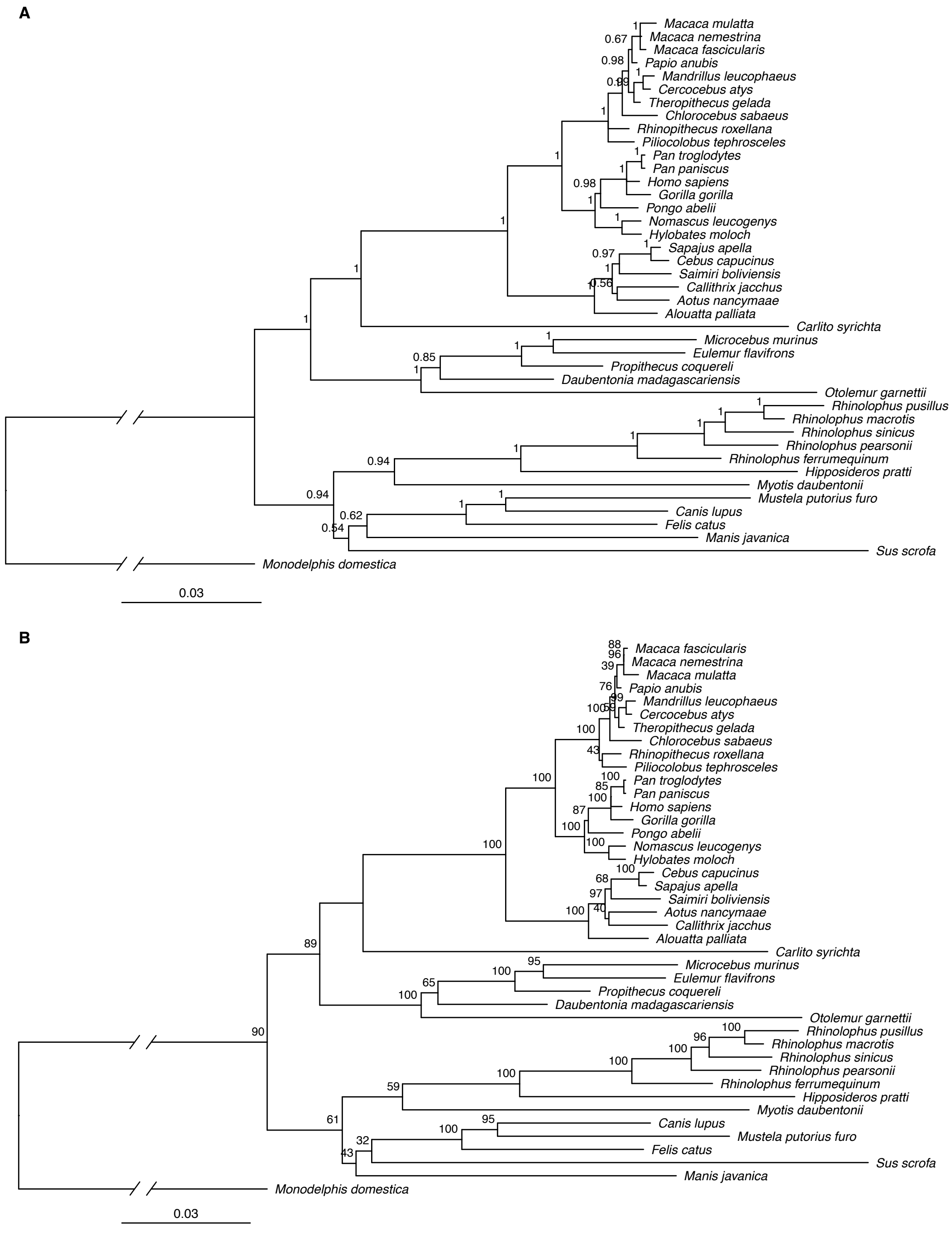


**Supplemental Figure S1.** *ACE2* gene trees built with (A) a Bayesian approach (Mr Bayes[3]) and (B) a maximum likelihood approach (RAxML[4]), using *Monodelphis domestica* as an outgroup. Node labels indicate (A) posterior probabilities or (B) bootstrap support. Scale bars indicate substitutions per site.

**Supplementary Figure S2.** Full-length alignment of ACE2 protein sequences.

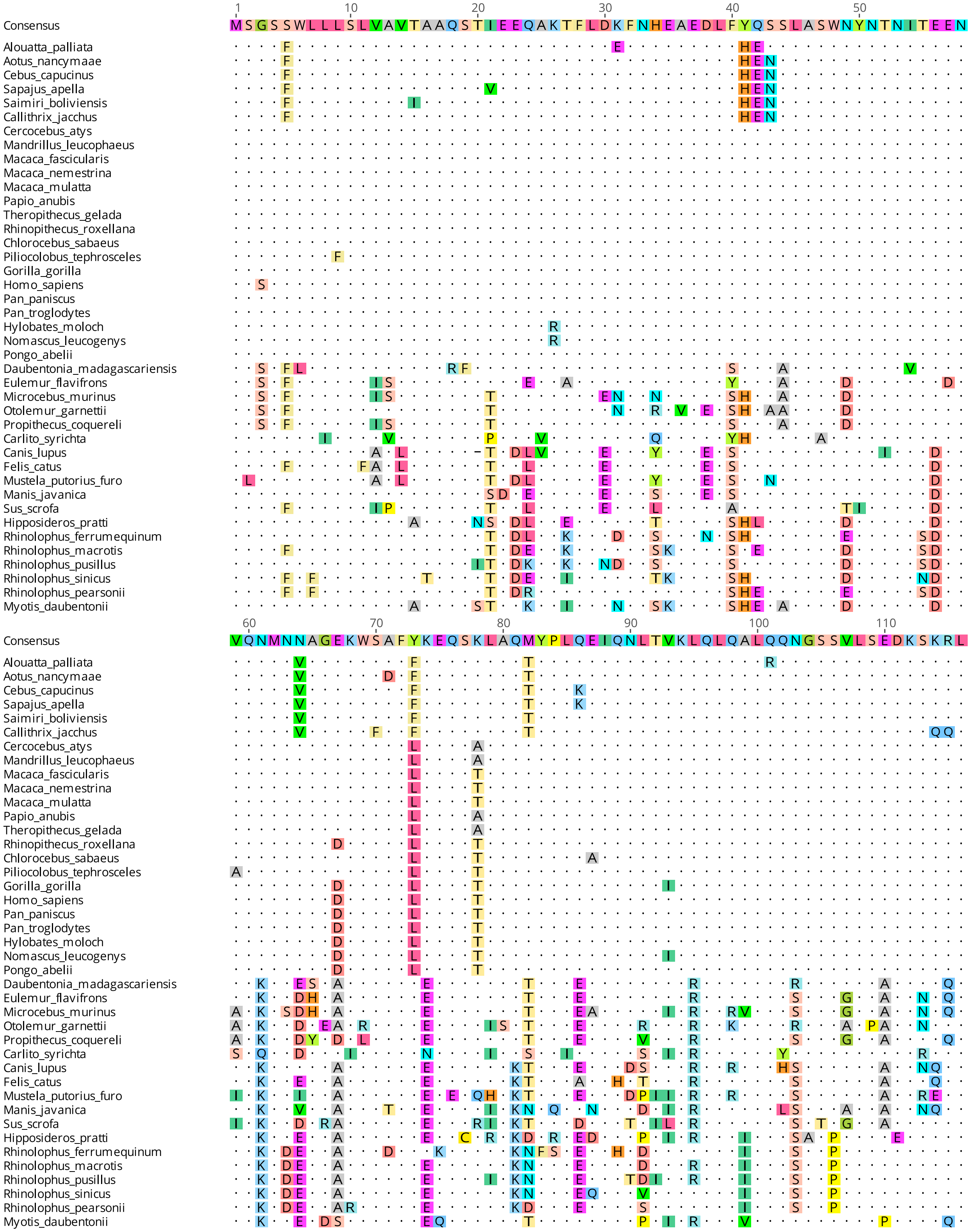

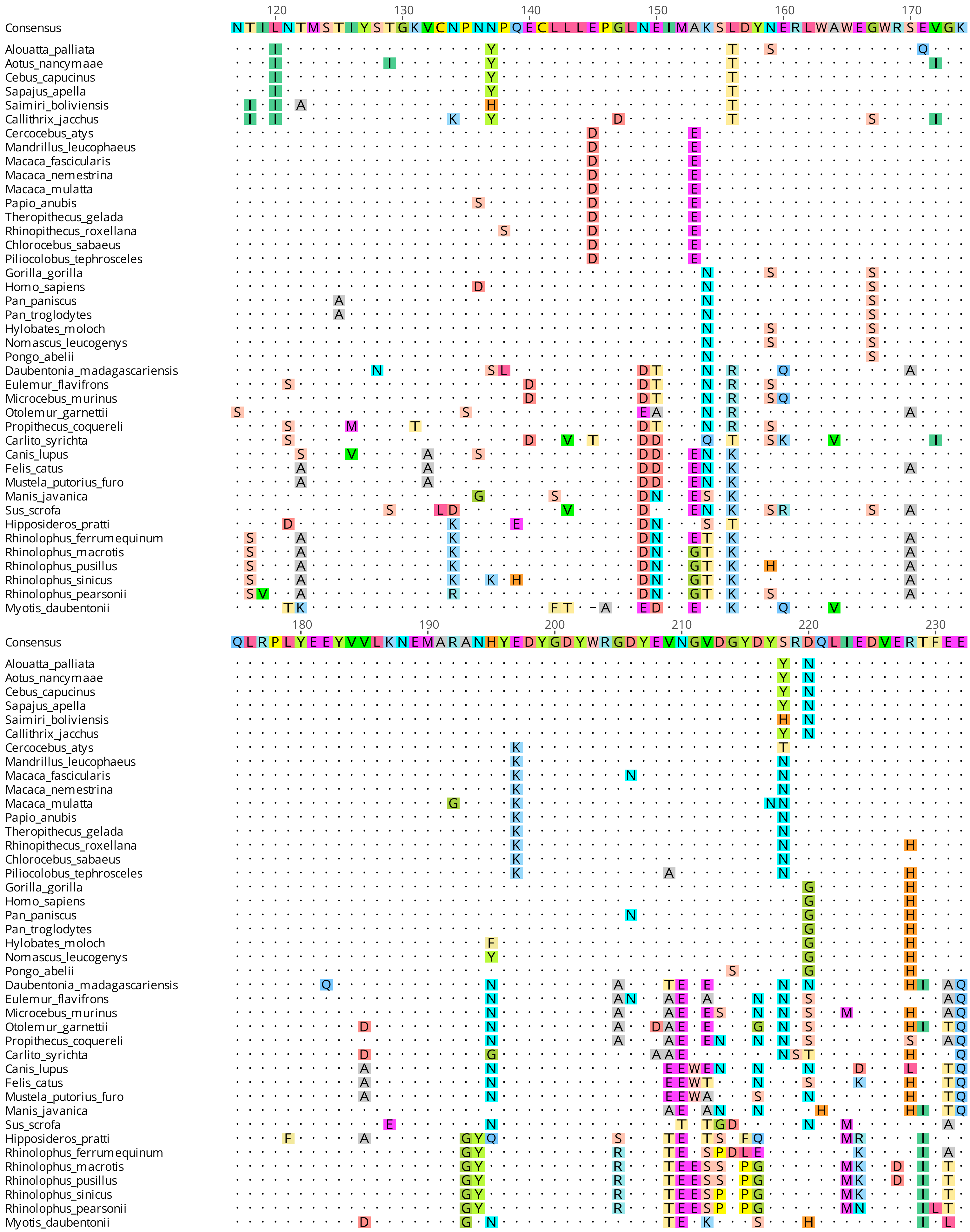
120 130 140 150 160 170
Consensus NTILNTMSTIYSTGKVCNPNNPQECLLEPGLNEIMAKSLDYNERLWAWEGWRSEVGK
180 190 200 210 220 230
Consensus QLRPLYEEYVVLKNEMARANHYEDYGDYWRGDYEVNGVDGYDYSRDQLIEDVERTFEE

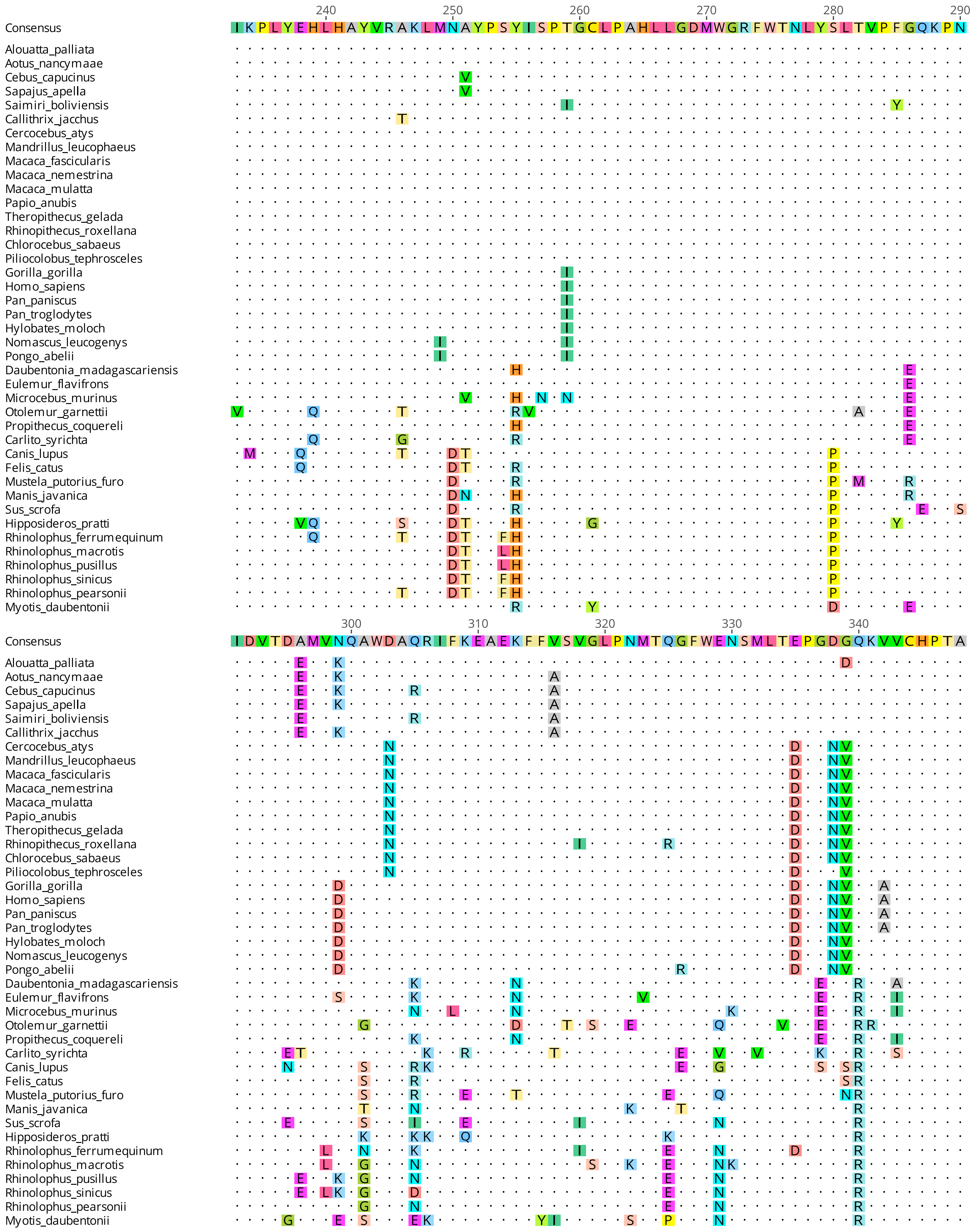

240
250
260
270
280
290
Consensus
IKPLYEHLHAYVRAKLMNAYPSYISPTGCLPAHLLGDMWGRFWTNLYSLTVPFGQKPN
300
310
320
330
340
IDVTDAMVNQAWDAQRIFKEAEKFFVSVGLPNMTQGFWENSMLTEPGDGQKVVCHPTA
Alouatta_palliata
Aotus_nancymaae
Cebus_capucinus
Sapajus_apella
Saimiri_boliviensis
Callithrix_jacchus
Cercocebus_atys
Mandrillus_leucophaeus
Macaca_fascicularis
Macaca_nemestrina
Macaca_mulatta
Papio_anubis
Theropithecus_gelada
Rhinopithecus_roxellana
Chlorocebus_sabaeus
Piliocolobus_tephrosceles
Gorilla_gorilla
Homo_sapiens
Pan_paniscus
Pan_troglodytes
Hylobates_moloch
Nomascus_leucogenys
Pongo_abelii
Daubentonia_madagascariensis
Eulemur_flavifrons
Microcebus_murinus
Otolemur_garnettii
Propithecus_coquereli
Carlito_syrichta
Canis_lupus
Felis_catus
Mustela_putorius_furo
Manis_javanica
Sus_scrofa
Hipposideros_pratti
Rhinolophus_ferrumequinum
Rhinolophus_macrotis
Rhinolophus_pusillus
Rhinolophus_sinicus
Rhinolophus_pearsonii
Myotis_daubentonii

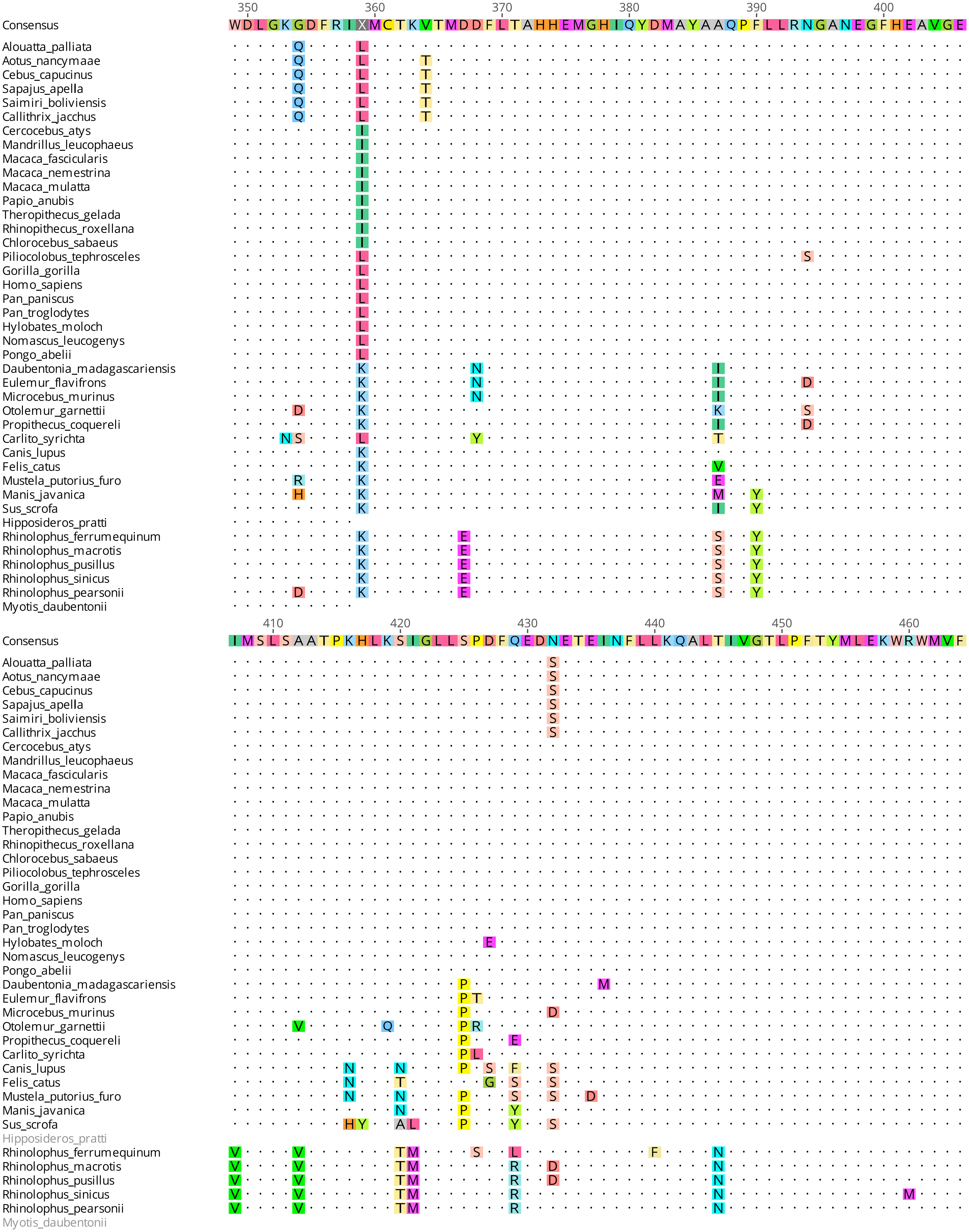
350 360 370 380 390 400
Consensus
WDLGKGDFRIXMCTKVTMDDFLTAHHEMGHIQYDMAYAAQPFLLRNGANEGFHEAVGE
Alouatta_palliata
Aotus_nancymaae
Cebus_capucinus
Sapajus_apella
Saimiri_boliviensis
Callithrix_jacchus
Cercocebus_atys
Mandrillus_leucophaeus
Macaca_fascicularis
Macaca_nemestrina
Macaca_mulatta
Papio_anubis
Theropithecus_gelada
Rhinopithecus_roxellana
Chlorocebus_sabaeus
Piliocolobus_tephrosceles
Gorilla_gorilla
Homo_sapiens
Pan_paniscus
Pan_troglodytes
Hylobates_moloch
Nomascus_leucogenys
Pongo_abelii
Daubentonia_madagascariensis
Eulemur_flavifrons
Microcebus_murinus
Otolemur_garnettii
Propithecus_coquereli
Carlito_syrichta
Canis_lupus
Felis_catus
Mustela_putorius_furo
Manis_javanica
Sus_scrofa
Hipposideros_pratti
Rhinolophus_ferrumequinum
Rhinolophus_macrotis
Rhinolophus_pusillus
Rhinolophus_sinicus
Rhinolophus_pearsonii
Myotis_daubentonii
410 420 430 440 450 460
Consensus
IMSLSAATPKHLKSIGLLSPDFQEDNETEINFLLKQALTIVGTLPFTYMLEKWRWMVF
Alouatta_palliata
Aotus_nancymaae
Cebus_capucinus
Sapajus_apella
Saimiri_boliviensis
Callithrix_jacchus
Cercocebus_atys
Mandrillus_leucophaeus
Macaca_fascicularis
Macaca_nemestrina
Macaca_mulatta
Papio_anubis
Theropithecus_gelada
Rhinopithecus_roxellana
Chlorocebus_sabaeus
Piliocolobus_tephrosceles
Gorilla_gorilla
Homo_sapiens
Pan_paniscus
Pan_troglodytes
Hylobates_moloch
Nomascus_leucogenys
Pongo_abelii
Daubentonia_madagascariensis
Eulemur_flavifrons
Microcebus_murinus
Otolemur_garnettii
Propithecus_coquereli
Carlito_syrichta
Canis_lupus
Felis_catus
Mustela_putorius_furo
Manis_javanica
Sus_scrofa
Hipposideros_pratti
Rhinolophus_ferrumequinum
Rhinolophus_macrotis
Rhinolophus_pusillus
Rhinolophus_sinicus
Rhinolophus_pearsonii
Myotis_daubentonii

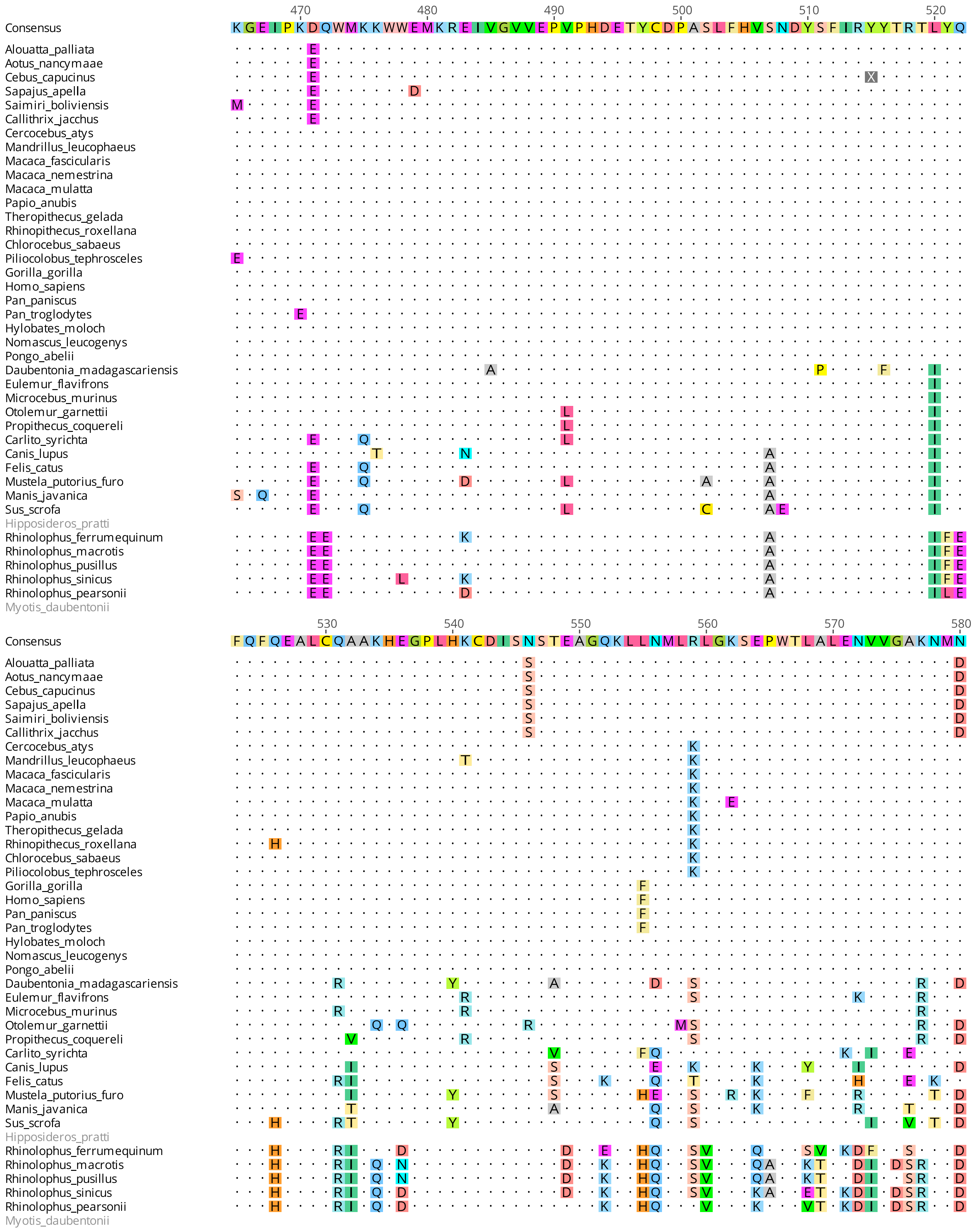
470 480 490 500 510 520
Consensus KGEIPKDQWMKKWWEMKREIVGVVEPVPHDETYCDPASLFHVSNDYSFIRYYTRTLYQ
530 540 550 560 570 580
Consensus FQFQEALCQAAKHEGPLHKCDISNSTEAGQKLLNMLRLGKSEPWTLALENVVGAKNMN
Alouatta_palliata
Aotus_nancymaae
Cebus_capucinus
Sapajus_apella
Saimiri_boliviensis
Callithrix_jacchus
Cercocebus_atys
Mandrillus_leucophaeus
Macaca_fascicularis
Macaca_nemestrina
Macaca_mulatta
Papio_anubis
Theropithecus_gelada
Rhinopithecus_roxellana
Chlorocebus_sabaeus
Piliocolobus_tephrosceles
Gorilla_gorilla
Homo_sapiens
Pan_paniscus
Pan_troglodytes
Hylobates_moloch
Nomascus_leucogenys
Pongo_abelii
Daubentonia_madagascariensis
Eulemur_flavifrons
Microcebus_murinus
Otolemur_garnettii
Propithecus_coquereli
Carlito_syrichta
Canis_lupus
Felis_catus
Mustela_putorius_furo
Manis_javanica
Sus_scrofa
Hipposideros_pratti
Rhinolophus_ferrumequinum
Rhinolophus_macrotis
Rhinolophus_pusillus
Rhinolophus_sinicus
Rhinolophus_pearsonii
Myotis_daubentonii

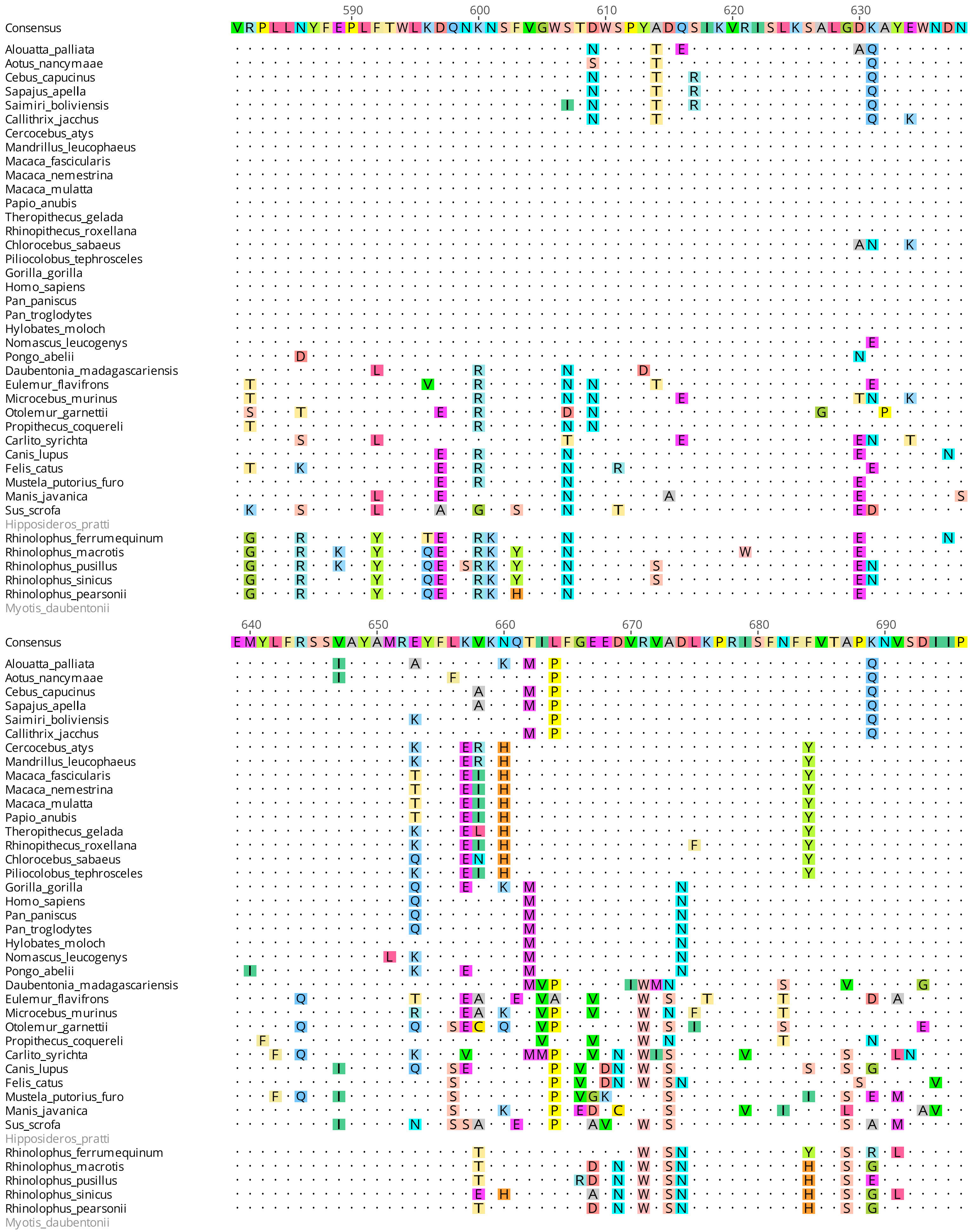
590 600 610 620 630
Consensus VRPLLNYFEPLFTWLKDQNKNSFVGWSTDWSPYADQSIKVRISLKSALGDKAYEWNDN
640 650 660 670 680 690
Consensus EMYLFRSSVAYAMREYFLKVKNQTILFGEEDVRVADLKPRISFNFFVTAPKNVSDIIP
Alouatta_palliata
Aotus_nancymaae
Cebus_capucinus
Sapajus_apella
Saimiri_boliviensis
Callithrix_jacchus
Cercocebus_atys
Mandrillus_leucophaeus
Macaca_fascicularis
Macaca_nemestrina
Macaca_mulatta
Papio_anubis
Theropithecus_gelada
Rhinopithecus_roxellana
Chlorocebus_sabaeus
Piliocolobus_tephrosceles
Gorilla_gorilla
Homo_sapiens
Pan_paniscus
Pan_troglodytes
Hylobates_moloch
Nomascus_leucogenys
Pongo_abelii
Daubentonia_madagascariensis
Eulemur_flavifrons
Microcebus_murinus
Otolemur_garnettii
Propithecus_coquereli
Carlito_syrichta
Canis_lupus
Felis_catus
Mustela_putorius_furo
Manis_javanica
Sus_scrofa
Hipposideros_pratti
Rhinolophus_ferrumequinum
Rhinolophus_macrotis
Rhinolophus_pusillus
Rhinolophus_sinicus
Rhinolophus_pearsonii
Myotis_daubentonii

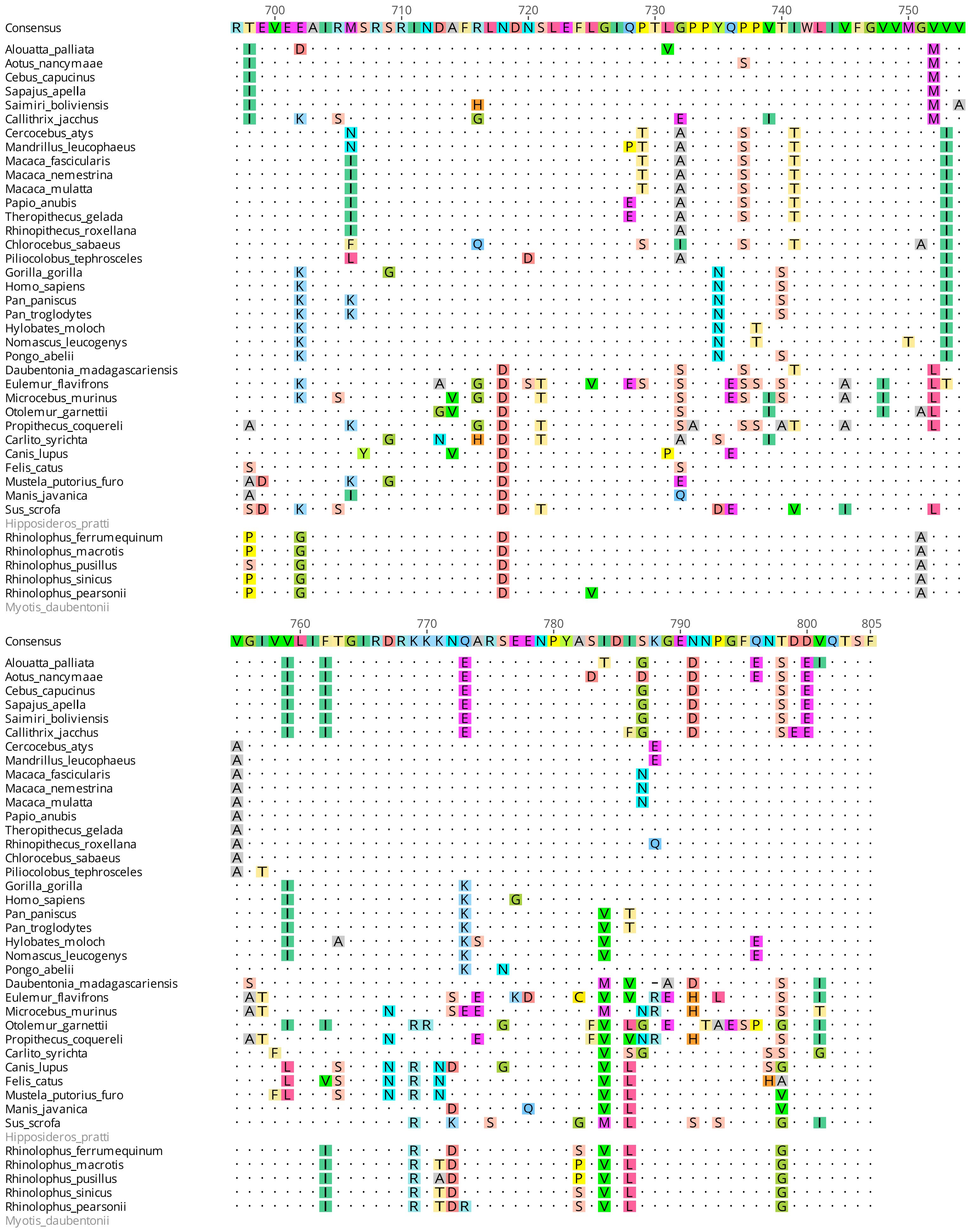
700 710 720 730 740 750
Consensus
RTEVEEAIRMSRSRINDAFRLNDNSLEFLGIQPTLGPPYQPPVTIWLIVFGVVMGVVV
760 770 780 790 800 805
VGIVVLIFTGIRDRKKKNQARSEENPYASIDISKGENNPGFQNTDDVQTSF
Alouatta_palliata
Aotus_nancymaae
Cebus_capucinus
Sapajus_apella
Saimiri_boliviensis
Callithrix_jacchus
Cercocebus_atys
Mandrillus_leucophaeus
Macaca_fascicularis
Macaca_nemestrina
Macaca_mulatta
Papio_anubis
Theropithecus_gelada
Rhinopithecus_roxellana
Chlorocebus_sabaeus
Piliocolobus_tephrosceles
Gorilla_gorilla
Homo_sapiens
Pan_paniscus
Pan_troglodytes
Hylobates_moloch
Nomascus_leucogenys
Pongo_abelii
Daubentonia_madagascariensis
Eulemur_flavifrons
Microcebus_murinus
Otolemur_garnettii
Propithecus_coquereli
Carlito_syrichta
Canis_lupus
Felis_catus
Mustela_putorius_furo
Manis_javanica
Sus_scrofa
Hipposideros_pratti
Rhinolophus_ferrumequinum
Rhinolophus_macrotis
Rhinolophus_pusillus
Rhinolophus_sinicus
Rhinolophus_pearsonii
Myotis_daubentonii
